# A cell atlas of chromatin accessibility across 25 adult human tissues

**DOI:** 10.1101/2021.02.17.431699

**Authors:** Kai Zhang, James D. Hocker, Michael Miller, Xiaomeng Hou, Joshua Chiou, Olivier B. Poirion, Yunjiang Qiu, Yang E. Li, Kyle J. Gaulton, Allen Wang, Sebastian Preissl, Bing Ren

## Abstract

Current catalogs of regulatory sequences in the human genome are still incomplete and lack cell type resolution. To profile the activity of human gene regulatory elements in diverse cell types and tissues in the human body, we applied single cell chromatin accessibility assays to 25 distinct human tissue types from multiple donors. The resulting chromatin maps comprising ∼500,000 nuclei revealed the status of open chromatin for over 750,000 candidate *cis*-regulatory elements (cCREs) in 54 distinct cell types. We further delineated cell type-specific and tissue-context dependent gene regulatory programs, and developmental stage specificity by comparing with a recent human fetal chromatin accessibility atlas. We finally used these chromatin maps to interpret the noncoding variants associated with complex human traits and diseases. This rich resource provides a foundation for the analysis of gene regulatory programs in human cell types across tissues and organ systems.

## INTRODUCTION

The human body is comprised of various organs, tissues and cell types, each with highly specialized functions. The genes expressed in each tissue and cell type – and in turn their physiologic roles in the body – are regulated by *cis*-regulatory elements such as enhancers and promoters (Carter and Zhao, 2020). These sequences dictate the expression patterns of target genes by recruiting sequence specific transcription factors (TFs) in a cell-type specific manner (Shlyueva et al., 2014). Upon binding of TFs, the regulatory elements frequently adopt conformational changes such that they are more accessible to endonucleases or transposases, enabling genome-wide discovery by combining with high throughput sequencing (Buenrostro et al., 2013; John et al., 2013; Klemm et al., 2019). However, conventional assays have, in large part, used heterogeneous tissues as input materials to produce population average measurements, and consequently, the current catalogs of candidate regulatory sequences in the human genome (Andersson et al., 2014; Meuleman et al., 2020; Moore et al., 2020; Roadmap Epigenomics et al., 2015; Shen et al., 2012) lack the information about cell type-specific activities of each element. This limitation has hampered our ability to study gene regulatory programs in distinct human cell types and to interpret the noncoding DNA in the human genome.

Genome wide association studies (GWAS) have identified hundreds of thousands of genetic variants associated with a broad spectrum of human traits and diseases. The large majority of these variants are non-coding (Claussnitzer et al., 2020). Observations that annotated *cis-*regulatory elements in disease-relevant tissues and cell types are enriched for non-coding risk variants (Ernst et al., 2011; Maurano et al., 2012; Roadmap Epigenomics et al., 2015) led to the hypothesis that a major mechanism by which noncoding variants influence disease risk is by altering transcriptional regulatory elements in specific cell types. However, annotation of these non-coding risk variants has been hindered by a lack of cell type-resolved maps of regulatory elements in the human genome. Whereas innovative approaches to distinguish causal variants from local variants in linkage disequilibrium (LD) using fine mapping (Wakefield, 2009), and to link variants to target genes using co-accessibility of open chromatin regions in single cells (Pliner et al., 2018) or 3-dimensional chromosomal contact-based linkage scores (Nasser et al., 2020), have made important strides toward the prioritization of causal variants and the prediction of their target genes, the annotation of candidate *cis-*regulatory elements (cCREs) in discrete human cell types has posed a longstanding technical challenge.

Single cell omics technologies, enabled by droplet-based, combinatorial barcoding or other approaches, have now enabled the profiling of transcriptome, epigenome and chromatin organization from complex tissues at single cell resolution (Grosselin et al., 2019; Klein et al., 2015; Lake et al., 2018; Luo et al., 2017a; Macosko et al., 2015; Preissl et al., 2018). In particular, combinatorial cellular barcoding-based assays such as single nucleus ATAC-seq (also known as sci-ATAC-seq (Cusanovich et al., 2015)) have permitted the identification of cCREs in single nuclei without the need for physical purification of individual cell types. The resulting data can be used to deconvolute cell types from mixed cell populations and to dissect cell type-specific transcriptomic and epigenomic states in primary tissues. While these tools have been applied to mammalian tissues including murine biosamples (Cusanovich et al., 2018; Lareau et al., 2019; Li et al., 2020; Preissl et al., 2018; Sinnamon et al., 2019), human fetal tissues (Domcke et al., 2020), and individual adult human organ systems (Chiou et al., 2019; Corces et al., 2020; Hocker et al., 2020; Wang et al., 2020), we still lack comprehensive maps of cCREs in the cell types comprising primary tissues of the adult human body.

In the present study we used a modified single-cell combinatorial indexing ATAC-seq (sci-ATAC-seq) protocol optimized for flash frozen primary tissues (Hocker et al., 2020; Preissl et al., 2018) to profile chromatin accessibility in 25 adult human tissue types from multiple donors. We profiled 472,373 nuclei from these tissues, grouped them into 54 cell types based on similarity in chromatin landscapes, and identified a union of 756,414 open chromatin regions and candidate CREs (cCREs) from the resulting maps. We then delineated gene regulatory programs in different human cell types, decomposed previous bulk chromatin accessibility maps, and characterized adult specific elements in different tissues and organ systems. Finally, we used the new cCRE atlas to interpret noncoding variants associated with complex human traits and diseases, demonstrating its utility in improving our understanding of polygenic human traits and revealing clinically relevant therapeutic targets for complex diseases. We created an interactive web atlas to disseminate this resource [CATLAS, Cis-element ATLAS] http://catlas.org/humantissue.

## RESULTS

### Single cell chromatin accessibility analysis of adult human primary tissues

In order to generate a cell type-resolved atlas of cCREs in the adult human body, we performed sci-ATAC-seq (Cusanovich et al., 2015; Preissl et al., 2018) with 70 primary tissue samples collected from 25 distinct anatomic sites in four postmortem adult human donors (Figure 1A, Table S1). Tissue samples were chosen to survey a breadth of human organ systems, including nine tissue types from across the gastrointestinal tract, four tissue types from the heart and peripheral vasculature, four female reproductive tissue types, three different endocrine tissue types, two tissue types from the integumentary system, and single tissue types from the muscular, peripheral nervous, and respiratory systems. Isolation of intact nuclei from these diverse primary tissue types, which differed in their nuclear compositions and sensitivities to mechanical dissociation, presented a technical challenge. We thus optimized nuclear isolation methods and buffer conditions for each tissue type (Table S2, see Methods). Subsequently, we generated sci-ATAC-seq datasets using a semi-automated workflow (Hocker et al., 2020; Li et al., 2020; Preissl et al., 2018) and sequenced resulting libraries to 7,651 raw sequence reads per nucleus on average, with a median read duplication rate of 44% (Table S3). Open chromatin fragments from these libraries were computationally assigned to individual nuclei using nucleus-specific DNA barcodes. We next filtered the single nucleus profiles based on stringent quality control criteria including an enrichment of reads at transcription start sites (TSS enrichment; TSSe) greater than 7-fold, and a minimum of > 1,000 mapped chromatin fragments per nucleus. Nuclei were further filtered for potential doublets, instances of 2 or more nuclei sharing a common barcode, using a version of Scrublet (Wolock et al., 2019) modified for sci-ATAC-seq (see Methods). Ultimately, we obtained high quality open chromatin profiles for 472,373 nuclei, with a median of 3,071 unique open chromatin fragments per nucleus and an average TSSe of 13.6 ± 4.5 per nucleus (Figure 1B, Figure S1, Table S3).

**Figure 1.**
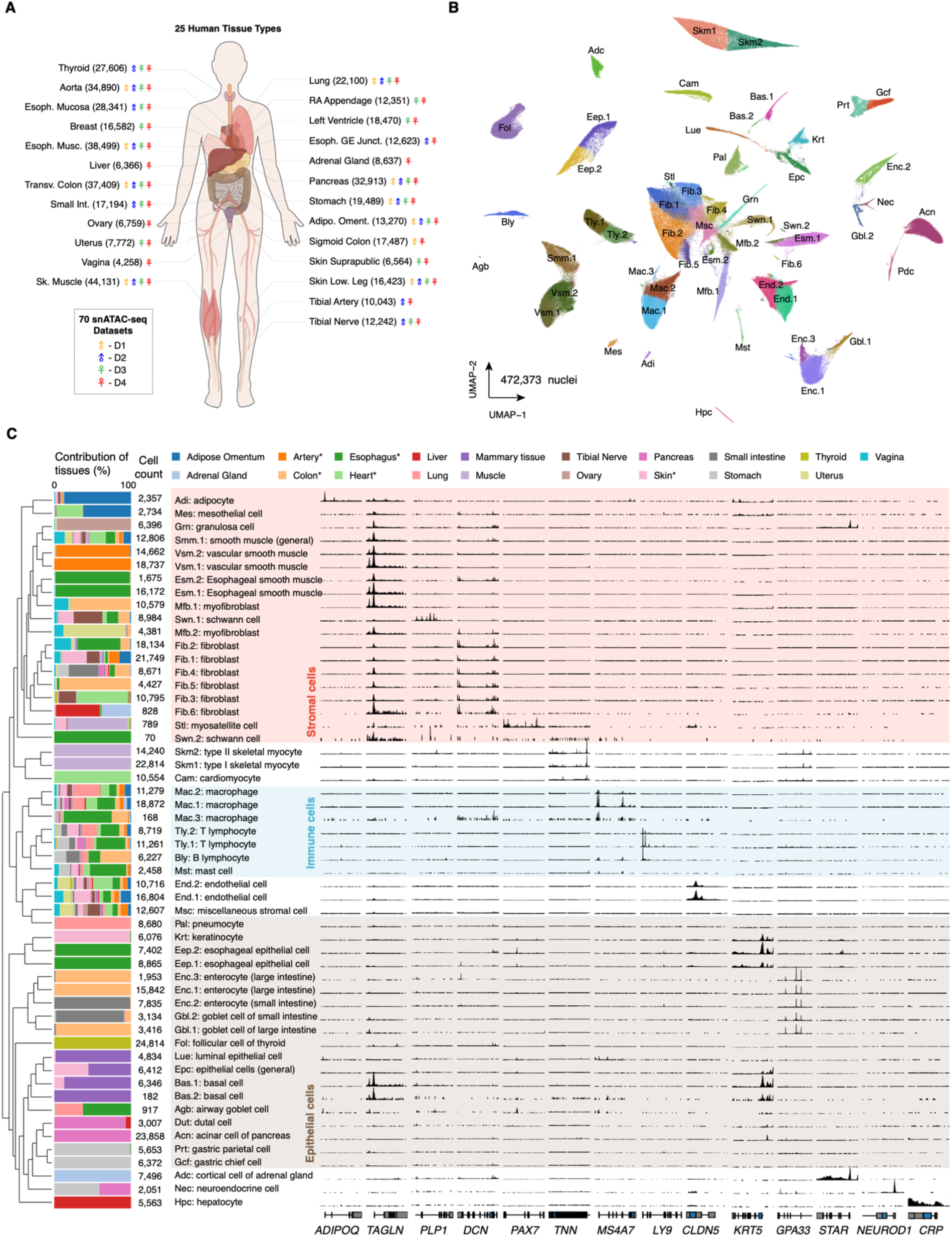
Single cell chromatin accessibility analysis of 25 adult human primary tissues. **A)** Overview of the study design. A total of 70 biosamples, representing 25 tissue types and obtained from up to four donors (D1 to D4), were used for sci-ATAC-seq assays. The number of nuclei profiled in each tissue was denoted in the parenthesis, along with the donor labels. **B)** Clustering of 472,373 nuclei identifying 54 distinct cell types. The visualization was generated using Uniform Manifold Approximation and Projection (UMAP) embedding. Clusters were annotated based on accessibility at promoters of marker genes as explained in the main text. Each dot in the scatter plot represents a nucleus. Nuclei are colored and labeled by cell type ID. The full names of the abbreviated cell type IDs are listed in panel **C**. **C)** Distribution of cell types across human tissues. The dendrogram on the left was created by hierarchical clustering of cell clusters based on chromatin accessibility. The bar chart represents relative contributions of tissues to cell clusters. * indicates categories representing multiple samples originating from similar tissues. Genome browser tracks on the right show aggregate chromatin accessibility profiles for each cell cluster at selected marker gene loci which were used for annotation.

Analyzing large and sparse single cell chromatin accessibility datasets has been challenging. According to a recent assessment of 10 popular computational methods for analyzing single cell ATAC-seq data (Chen et al., 2019), SnapATAC was the only method able to cluster > 80,000 cells without sacrificing accuracy. In the latest development of SnapATAC, we utilized the Nyström method (Bouneffouf and Birol, 2016) to further improve the scalability of the algorithm to handle millions of cells, an indispensable feature for atlas-scale studies. When dealing with samples from diverse biological backgrounds, individual and batch effects are inevitable and pose further challenges to integrative analysis. We built upon the Mutual Nearest Neighbor batch-effect-correction method (Haghverdi et al., 2018) to develop a variant called Iterative Mutual Nearest Centroid algorithm to correct for donor or batch effects with added scalability and flexibility (Figure S2A-C, see Methods). After dimensionality reduction and batch correction, we applied the Leiden algorithm (Traag et al., 2019) to identify cell clusters. To determine the optimal number of cell types present in the dataset, we surveyed the stability of clustering results upon simulated perturbation under different parameters (Figure S2D, see Methods). This analysis yielded a total of 54 distinct clusters with high reproducibility and diversity (Figure 1B, Figure S2C-D, Table S4).

### Annotation of major and sub-classes of human cell types

To annotate the resulting cell clusters, we first curated a set of marker genes from the PanglaoDB single cell RNA-seq marker gene database (Franzén et al., 2019) corresponding to expected human cell types. We utilized chromatin accessibility at the promoter, defined as ±1000 bp relative to transcription start sites (TSS), as a proxy for gene activity and computed cell-type enrichment scores for each of the 54 clusters, and created initial cell cluster annotations based on these cell-type enrichment scores (Figure S3A, see Methods). We next manually reviewed these assignments and made adjustments based on focused consideration of marker gene accessibility. Reassuringly, enrichment of Gene Ontology (GO) terms for genes linked to restricted peaks in a given cluster was in agreement with presumed functions of assigned cell types (Figure S4). Finally, we compared our single-cell chromatin accessibility atlas with a recent single cell transcriptional atlas of adult human tissues (Han et al., 2020). Correlating promoter accessibility profiles from sci-ATAC-seq clusters with gene expression profiles from scRNA-seq clusters, we found that the cell types with the highest correlation across datasets were concordantly annotated in the majority of cases (Figure S3B). Altogether, we were able to annotate 53 of the 54 clusters (98%) with a cell type label (Table S5). For example, we annotated three macrophage clusters based on accessibility at marker genes including *MS4A7 (Gingras et al., 2001)*, and one adipocyte cluster based on accessibility at *ADIPOQ (Hu et al., 1996)* (Figure 1C). Encouragingly, prevalent cell types detected in a majority of tissue samples including macrophages, lymphocytes, endothelial cells, and smooth muscle cells clustered based on cell type rather than tissue of origin or individual (Figure 1C, Table S4, Figure S5).

Most of these cell types were found to exhibit high tissue specificity. For example, some highly specialized cell types such as granulosa cells, follicular cells, parietal cells, chief cells, pneumocytes, keratinocytes, and hepatocytes were restricted to only one tissue type, reflecting their tissue-specific functions (Figure 1C, Table S4, Figure S5). We further annotated five clusters of lower gastrointestinal (GI) tract epithelial cells that could be classified as either enterocytes or goblet cells, but which were differentially clustered according to whether nuclei originated in the small intestine (Enc. 2, Gbl.2) or colon (Enc.1 & 3, Gbl.1; Figure 1C). On the other hand, tissue-resident fibroblasts unbiasedly clustered into six subtypes with diverse tissues of origin for each (Fib.1-6; Figure 1C). Our analysis also revealed rare cell types with distinct chromatin accessibility profiles such as mesothelial cells (0.58% of total nuclei) and satellite cells or muscle stem cells (0.17% of total nuclei). During annotation, we noticed that some cell clusters appeared to contain multiple closely related but distinct cell types. For example, the neuroendocrine cell cluster consisted of cells from both stomach and pancreas, likely representing a mixture of pancreas- and stomach-specific hormone-producing cells. To further dissect the heterogeneity within our identified cell clusters, we performed another round of clustering on cell clusters that contained at least 1,000 nuclei and showed minimal batch effects (see Methods). We were able to identify more than one subcluster in 15 out of 27 major cell classes satisfying the above criteria (Figure S6A). In particular, the neuroendocrine cell cluster was further divided into three clusters that could be annotated as beta cells, alpha cells, and gastric neuroendocrine cells based on accessibility at maker genes including *INS*, *GCG*, and *GHRL*, respectively (Chiou et al., 2019; Kojima et al., 1999) (Figure S6). Moving forward, we focused our subsequent analyses on the 54 cell clusters defined by our initial data-driven approach due to our high level of confidence in their stability, reproducibility, and cell type annotation.

### An atlas of cCREs in adult human cell types

We annotated cCREs in each of the 54 primary cell types defined above. To do so, we aggregated chromatin accessibility profiles from all nuclei comprising each cell cluster and identified open chromatin regions using the MACS2 software package (Zhang et al., 2008) (Figure 2A). We then merged peaks from all cell clusters to form a union set of 756,414 open chromatin regions and termed these as cCREs (Figure 2A-C, Table S6, Supplementary file with accessibility for each cCRE downloadable from http://catlas.org/humantissue). These cCREs covered 11.4% of the human genome, and 92.7% of them overlapped with previously annotated cCREs based on bulk DNase-seq and ChIP-seq assays of human tissues, cell lines, and primary cell biosamples by the ENCODE consortium (Meuleman et al., 2020; Moore et al., 2020) (Figure 2B). Genome-wide, cCREs located at transcription start sites or near promoter regions tended to have elevated chromatin accessibility, were less likely to vary between different cell types, and displayed higher levels of sequence conservation than gene-distal cCREs and genomic background (Figure 2D-E). By contrast, gene-distal cCREs tended to be more variable chromatin accessibility (Figure 2D), suggesting the presence of shared programs of highly accessible promoter-proximal cCREs alongside variable programs of gene-distal cCREs across human cell types.

**Figure 2.**
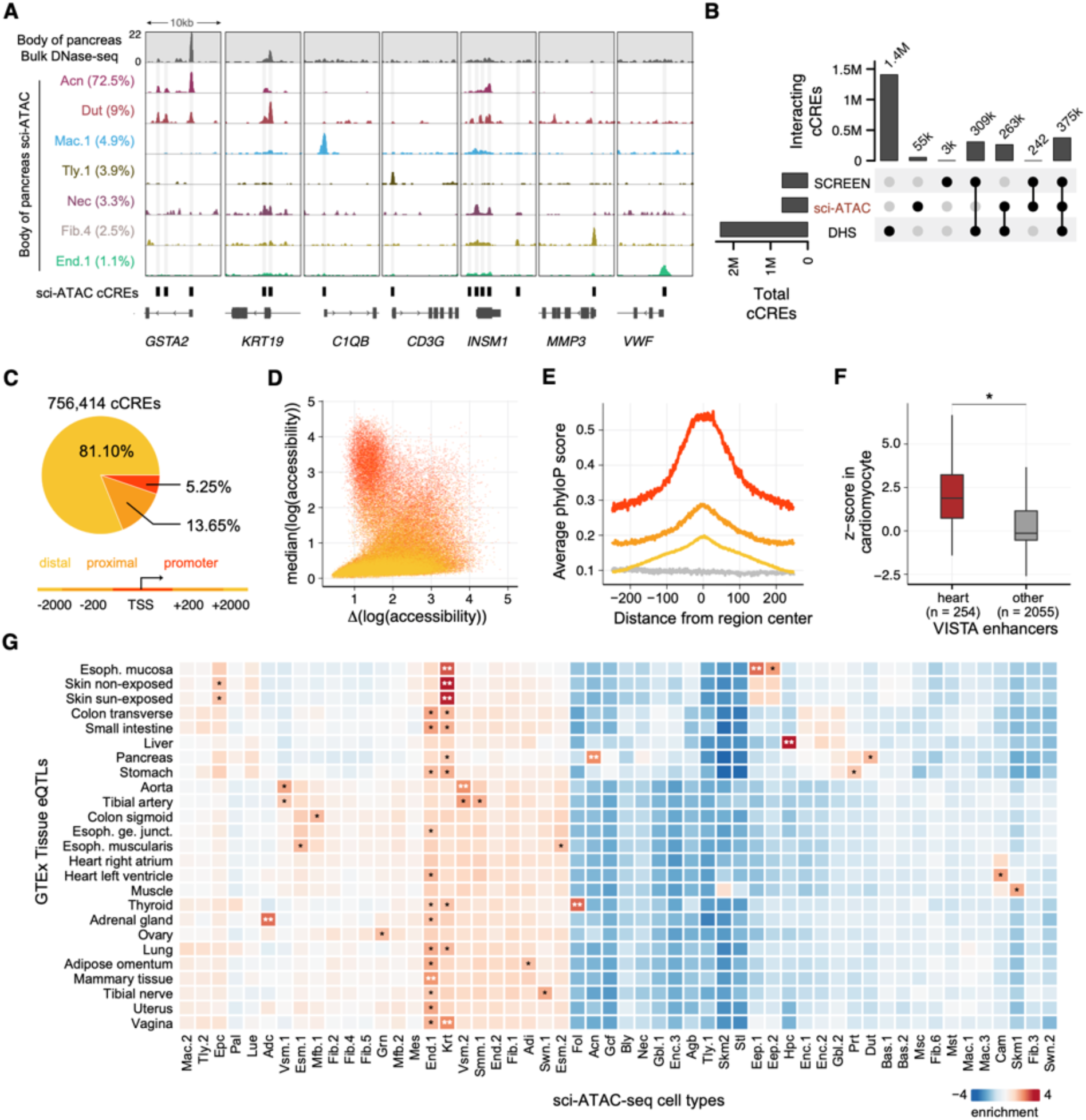
An atlas of cCREs in adult human cell types. **A)** Genome browser tracks comparing sci-ATAC-seq with bulk DNase-seq data from the ENCODE consortium (Accession: ENCSR464TKV) for detecting accessible regions in body of pancreas as an example of a complex heterogeneous tissue containing multiple cell types. **B)** Intersection between three cCRE catalogues showing that the majority of identified cCREs in the present study are supported by previous functional annotations released by the ENCODE consortium. **C)** Distribution of 756,414 cCREs across the human genome. Based on their distances to annotated gene transcription start sites, we classified cCREs into one of the three groups: promoter, promoter-proximal and distal. **D)** Scatter plot showing the three groups of cCREs based on median and range (difference between maximum and minimum) of chromatin accessibility across cell clusters. Each dot represents a cCRE, colored by groups in **C**. **E)** Average phyloP (Pollard et al., 2010) scores of cCREs stratified by groups defined in **c**. Genomic background is indicated in gray. **F)** Boxplot comparing validated heart-specific *in vivo* enhancers from VISTA database against other enhancers from VISTA database based on their chromatin accessibility in cardiomyocytes. **G)** Z-scores for enrichment of GTEX eQTLs from corresponding tissues in each cell cluster. *: p < 0.05, **: p < 0.01.

To assess the function of the above cCREs, we compared them with current catalogs of validated enhancers (Visel et al., 2007) and expression quantitative trait loci (eQTLs) - sequence variants that are statistically correlated with changes in gene expression in a tissue-specific fashion (Consortium, 2020). We first compared our cCREs with the VISTA database (Visel et al., 2007), and found that they were enriched for enhancers validated in transgenic mice in a cell type-specific fashion (Figure 2F). We next asked whether our cCREs were enriched for eQTLs annotated by the GTEx Project in the 25 matching adult tissue types. We discovered cell type-specific enrichments for 24 out of 25 sets of tissue eQTLs (Figure 2G). As expected, tissue eQTLs were most strongly enriched within cCREs when the corresponding cell type comprised a large proportion of nuclei identified in the tissue (Figure S7). For example, thyroid tissue eQTLs were strongly enriched within cCREs annotated in follicular cells (p = 0.0024), which made up 90.4% of total nuclei from thyroid tissue. On the other hand, tissue eQTLs from heterogenous tissue types such as transverse colon tended to display weaker overall enrichment in cell type cCREs, as well as a tendency to be enriched within cCREs of prevalent cell types that could be identified in most primary tissues, such as endothelial cells (Figure 2G, Figure S7). Taken together, these results suggest that bulk tissue eQTLs best represent sequence variants associated with gene expression for abundant cell types and homogenous tissues, and may be less representative for rarer cell types within homogenous tissues or for unique cell types from heterogenous tissues.

### Delineation of cell-type specificity of human cCREs

Cell fate determination in part depends on the establishment of specific *cis*-regulatory programs modulating gene expression. To characterize the cell-type specificity of cCREs, we organized the 756,414 cCREs into 51 *cis*-regulatory modules (CRMs), with elements in each CRM sharing similar chromatin accessibility patterns across all the cell types defined in the current study (Figure 3A, see Methods). We further annotated candidate functions of CRMs based on GREAT biological process ontology terms (McLean et al., 2010) (Figure 3B, Table S7). These analyses revealed that the majority of CRMs were limited either to single cell types or to groups of cell types that reflected cellular lineages. For example, one CRM related to the maintenance of gastrointestinal epithelium showed preferential accessibility in goblet cells (Module 8; Figure 3A-B), whereas two additional CRMs related to regulation of actin filament organization and glucose transport showed strong shared accessibility across all lower gastrointestinal epithelial cell types, including both goblet cells and enterocytes (Modules 9 and 10; Figure 3A-B). Broadly, CRM annotations reflected the physiologic functions of the cell types with which they were associated. For example, follicular cells were enriched for a CRM related to the regulation of iodide transport, hepatocytes for a CRM related to steroid metabolism, and skeletal myocytes for CRMs related to the regulation of muscle structure development (Modules 12, 14 and 34; Figure 3A-B).

**Figure 3.**
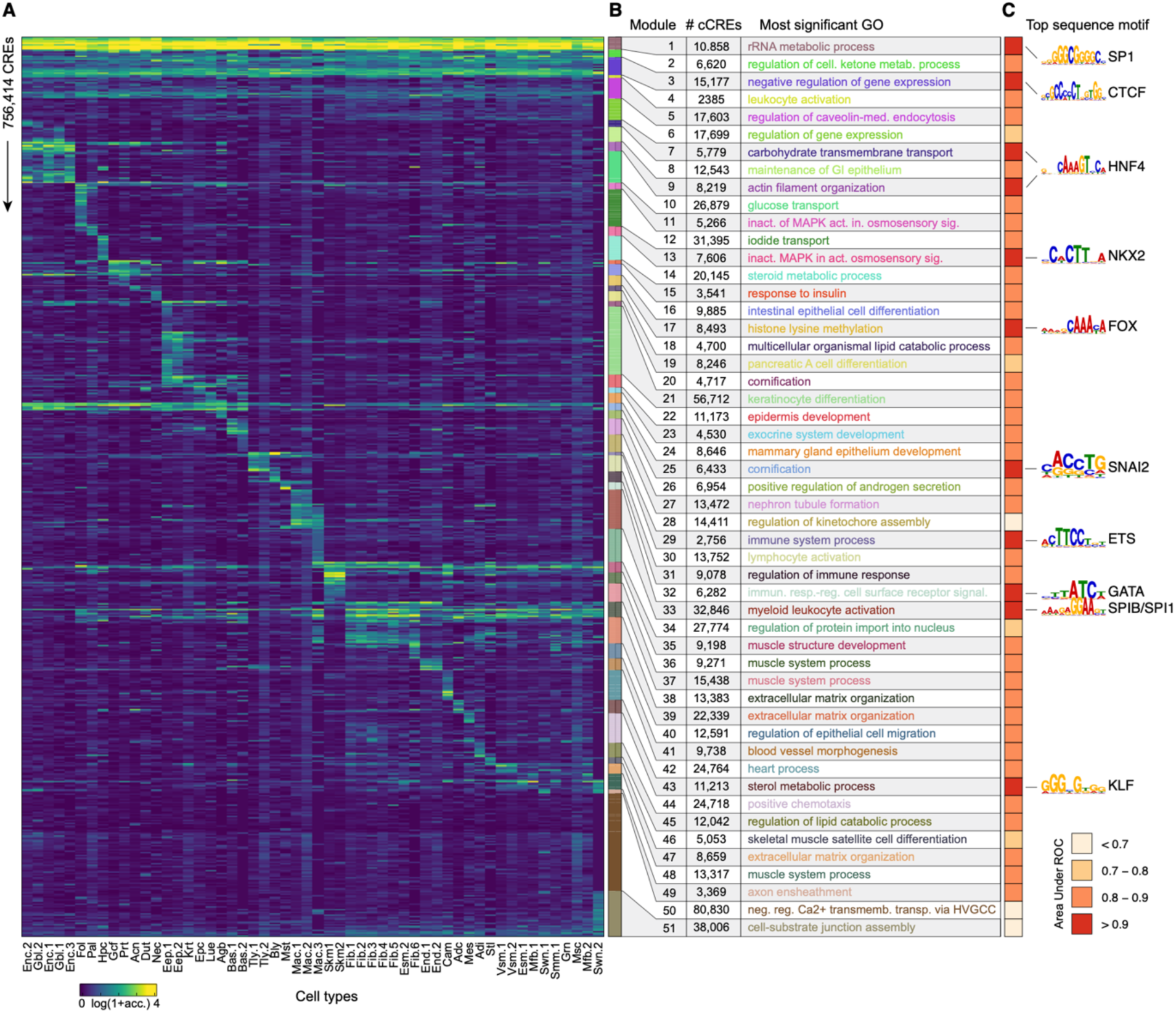
Delineation of cell-type specificity of human cCREs. **A)** Heatmap representation of chromatin accessibility for 756,414 cCREs across 54 human cell types. Each row represents an individual cCRE, while each column represents a cell type. The cell type ID is the same as Figure 1C. Color represents relative chromatin accessibility. cCREs were organized into 51 modules by clustering (see Methods). Color bars to the right depict the module ID. **B)** Top GREAT ontology enrichment (significance level: FDR < 0.01) for each cCRE module. **C)** Heatmap representation of area under the receiver operating characteristics (AUROC) across 51 cCRE modules. We trained a 51-class convolutional neural network to predict the module class for each cCRE using DNA sequences as the features (Figure S8). For each module the AUROC measures how well the classifier distinguishes cCREs belonging to the target module from the rest. On the right of the heatmap the top sequence motif features for the best performing modules are shown. Motifs were extracted from the neural network model using the TF-MoDISco algorithm (Shrikumar et al., 2018).

Cell type-specific *cis*-regulatory programs arise from combinatorial actions of sequence-specific TFs. To investigate the extent to which DNA sequence determined the cell type-specific accessibility patterns manifested in the 51 CRMs defined above, we trained a 51-class convolutional neural network using genomic sequence as the sole feature to predict module membership for each cCRE, and measured the area under the resulting ROC curve (AUROC) as a metric of classifier performance (Figure S8, see Methods). For 44 out of 51 modules, cCRE sequence alone could predict module membership with an AUROC > 0.80 (Figure 3C), suggesting that DNA sequence may play a pivotal role in forming diverse CRMs across cell types. To derive the sequence features that allowed our neural network to distinguish between cCRE modules, we applied the Transcription Factor Motif Discovery from Importance Scores (TF-MoDISco) software package, which deciphers consolidated motifs learned by DNA sequence-based neural networks (Shrikumar et al., 2018). Comparing these learned motifs with catalogued TF motifs (Weirauch et al., 2014) revealed module-specific TF motifs (Figure 3C). For example, sequence features matching the SP1 motif distinguished a module with strong accessibility in all identified cell types from other modules, consistent with the original description of SP1 as a regulator of ubiquitously-expressed housekeeping genes (Black et al., 2001) (Module 1; Figure 3C). Similarly, sequence features matching the NKX2 motif distinguished a module unique to pneumocytes, in line with the role of NKX2 in regulating the production of pulmonary surfactant (Bingle, 1997; Bohinski et al., 1994) (Module 13; Figure 3C). In addition to previously-characterized associations, we also report previously undefined TF associations with adult human cell types that are challenging to study in their *in vivo* tissue contexts: for example, the motif of the FOX TF family (Golson and Kaestner, 2016) differentiated modules accessible in gastric chief cells and parietal cells (Module 17; Figure 3C), and the motif of the KLF family (McConnell and Yang, 2010) differentiated a module accessible in adrenal cortical cells (Module 43; Figure 3C).

### Decomposition of bulk chromatin accessibility data using single cell chromatin atlas

Previous studies to assay chromatin accessibility have utilized biosamples including primary tissues, marker-isolated primary cells, cultured primary cells, *in vitro* differentiated cell lines, and immortalized cell lines (Kundaje et al., 2015; Meuleman et al., 2020; Moore et al., 2020; Stunnenberg et al., 2016). In order to quantify how closely these datasets from bulk assays resembled chromatin signatures from individual adult human cell types profiled in the current study, we compiled publicly available bulk ATAC-seq and DNase-seq datasets and measured their correlation with adult human cell type chromatin accessibility profiles from sci-ATAC-seq. Biosamples exhibited a wide range of correlation scores with human cell types. In aggregate however, primary cell type biosamples resembled adult cell types profiled in the current study more closely than did bulk tissue or cell line biosamples (Figure S9, Table S8).

Analysis of chromatin accessibility in bulk primary human cancer biosamples from The Cancer Genome Atlas (TCGA) (Cancer Genome Atlas Research et al., 2013) has been shown to be a powerful tool for the characterization of abnormal gene regulatory elements in cancer and the classification of tumor subtypes with prognostic importance (Corces et al., 2018), but previous analyses were performed on bulk tumor samples and lacked information about the cell types responsible for signature chromatin accessibility patterns. We thus used our cell atlas to deconvolute bulk chromatin accessibility datasets from human primary tumor biosamples (Corces et al., 2018) into non-tumor cell classes based on chromatin accessibility features. We developed a support vector regression (SVR) based method for deconvolution. We showed that our method performed well on a variety of benchmarking datasets (median coefficient of determination = 0.941, Figure S10A), and that the performance was robust against the choice of features, a wide range of sequence depths, and the introduction of artificial noise (Figure S10B-E, see Methods). We further benchmarked this approach by deconvoluting 21 bulk DNase-seq datasets from human stomach tissue, which revealed signatures of parietal cells across life stages but signatures of gastric chief cells only in child and adult timepoints, consistent with the histologic appearances of these cell types in the developing human stomach (Roy and Roy, 2016). We finally applied our deconvolution approach to 275 bulk ATAC-seq biosamples from 13 primary cancer types, and found that predicted cell type composition varied greatly between cancer types (Figure S10G). For example, whereas primary thyroid carcinomas (THCA), adrenocortical carcinomas (ACC), and liver hepatocellular carcinomas (LIHC) contained biosamples with dominant chromatin signatures from follicular cells, adrenal cortical cells, and hepatocytes respectively, primary stomach adenocarcinomas (STAD) contained a mixture of biosamples with chromatin signatures from immune cells, goblet cells, enterocytes, and parietal cells. Primary breast invasive carcinomas (BRCA) in particular showed a marked variety of cell type signatures, containing biosamples with chromatin signatures from mammary luminal epithelial cells, general epithelial cells, basal cells, airway goblet cells, and adipocytes (Figure S10H). Based on these chromatin signatures, breast cancer biosamples could be further categorized into cellular subtypes that corresponded with bulk gene expression patterns as well as prognostic features (Figure S10I-K).

### Identification and characterization of adult-specific human cCREs

We next compared adult cell type chromatin accessibility signatures with their corresponding fetal cell types in order to investigate life stage-specific chromatin signatures. Drawing from a recent cell atlas of chromatin accessibility in human fetal tissues (Domcke et al., 2020), we first selected fetal tissue types that matched those assayed in the current study and quantified correlations between fetal and adult cell types based on chromatin accessibility over a merged set of cCREs (see Methods). Out of 41 adult cell types from matching tissue types, 31 had chromatin signatures that were significantly correlated with at least one fetal cell type (Figure 4A). Interestingly, while some of these cell types such as cardiomyocytes, Schwann cells, and endothelial cells exhibited highly correlated chromatin signatures between fetal and adult stages (P < 0.01), other comparably specialized adult cell types, such as satellite cells and skeletal myocytes, were not significantly correlated with their fetal counterparts (Figure 4A). Comparing chromatin accessibility between fetal and adult stages genome-wide, we found a total of 208,024 adult-specific cCREs (Figure 4B).

**Figure 4.**
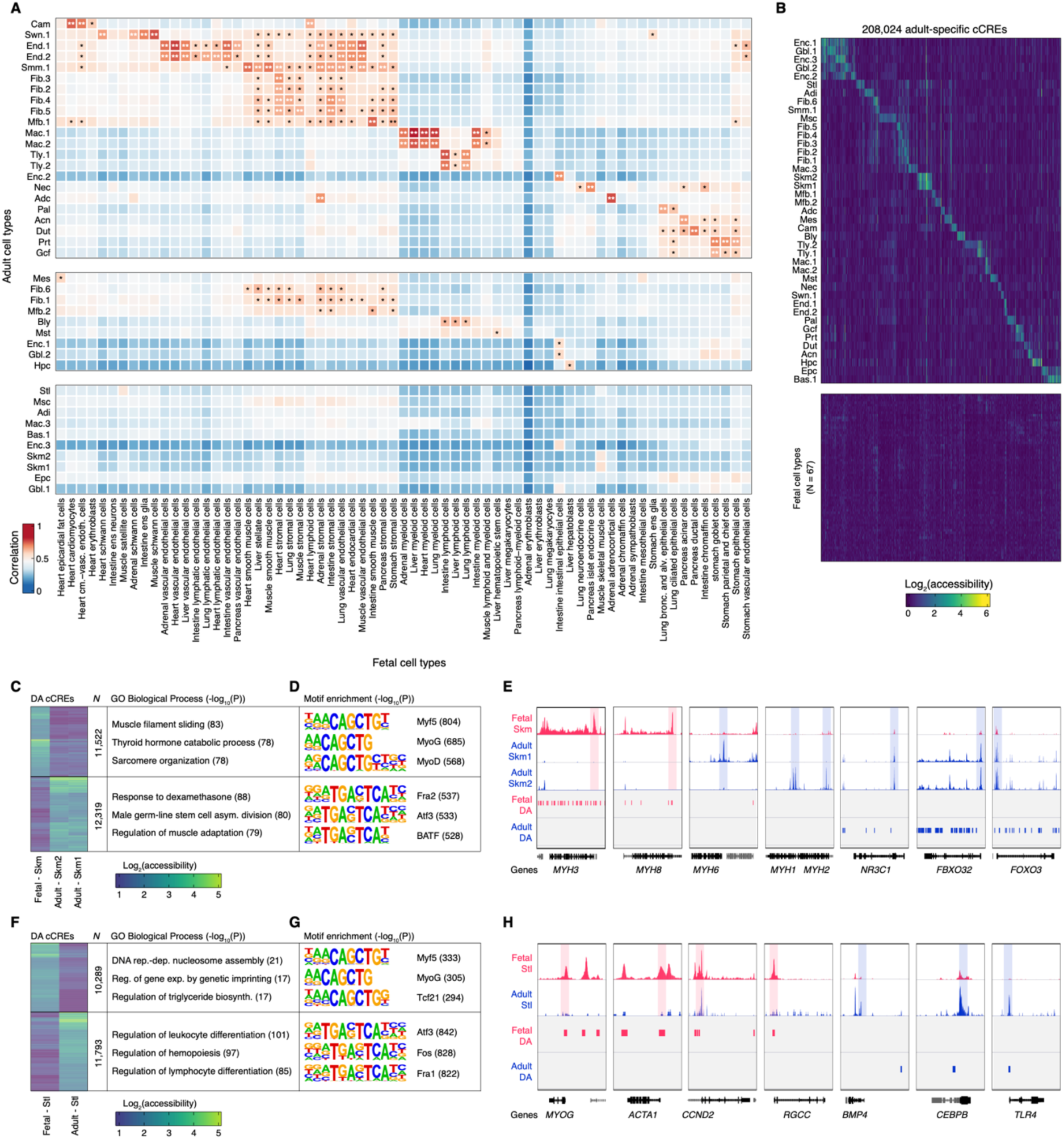
Comparison of chromatin accessibility between fetal and adult stages. **A)** Heatmap showing similarity between fetal (column) and adult (row) cell types in matching tissues. Color represents Pearson correlation coefficient. *: p < 0.05, **: p < 0.01. **B)** Heatmap representation of 208,024 adult-specific cCREs. Color represents log-transformed normalized signal. **C)** Heatmap representation of 23,841 differentially accessible (DA) cCREs between fetal skeletal myocytes and adult skeletal myocytes along with the top three GREAT biological process ontology enrichments (McLean et al., 2010) for adult and fetal skeletal myocyte DA cCREs. Color represents log-transformed normalized signal. **D)** Top three known TF motifs enriched within fetal and adult skeletal myocyte DA cCREs identified by HOMER (Heinz et al., 2010). **E)** Genome browser tracks showing chromatin accessibility for fetal and adult skeletal myocytes along with DA cCREs between the adult and fetal skeletal myocytes. Indicated genes are shown in black, other genes are shown in gray. Transcription start sites of the indicated genes are shaded in red and blue. **F-H** represent the same analyses performed in **C-D** for 22,082 DA cCREs between fetal satellite cells and adult satellite cells.

To uncover the gene regulatory programs that may underlie developmental functions, we next determined adult and fetal-specific cCREs in cell types that showed pronounced differences in chromatin accessibility between life stages. Skeletal myocytes, for example, differentiate substantially during pre and post-natal development (Chal and Pourquié, 2017) and showed poorer correlation between life stages than other human cell types (Figure 4A). In total, we identified 23,841 differentially accessible (DA) cCREs between fetal and adult skeletal myocytes (Figure 4C). DA cCREs in fetal myocytes were associated with biological processes such as muscle filament sliding and sarcomere organization, and were strongly enriched for motifs of myogenic regulatory TFs (MRFs) which orchestrate normal myogenesis (Mary Elizabeth Pownall et al., 2002), including myogenic factor 5 (Myf5), myogenin (MyoG), and myoblast determination factor (MyoD) (Figure 4C-D), highlighting the potential role of these elements in regulating myogenic processes and the expression of fetal-specific myosin isoforms. On the other hand, adult skeletal myocyte DA cCREs were associated with biological processes related to glucocorticoid response and regulation of muscle adaptation, and were enriched for the motifs of AP-1 complex members Fra2, Atf3, and BATF (Figure 4C-D), suggesting a potential role for these elements in regulating transcriptional responses to steroid hormones and adaptation to the differential contractile activity and loading conditions of adult skeletal muscle. In line with our ontology results and with established patterns of myosin isoform expression across the human lifespan (Schiaffino and Reggiani, 2011; Schiaffino et al., 2015; Stuart et al., 2016), we discovered DA cCREs at loci encoding marker genes of pre-natal myocytes including *MYH3* and *MYH8*, the heavy chains of embryonic and neonatal myosin respectively, as well as markers of type I (slow) and type II (fast) twitch adult myocytes including *MYH6* and *MYH1/MYH2* respectively (Figure 4E).

Encouraged by these findings, we next examined differences in chromatin accessibility between fetal and adult satellite cells or muscle stem cells (Yin et al., 2013), which similarly to skeletal myocytes were not significantly correlated between life stages (Figure 4A). Fetal satellite cells are highly proliferative and play an important role in the rapid expansion of skeletal muscle mass in the pre-natal period, whereas adult satellite cells represent a small pool of quiescent myocyte precursors (Chal and Pourquié, 2017). Thus, knowledge of the regulatory elements that modulate these processes could yield important insights into the regulation of muscle regeneration. Our analysis revealed 22,082 differentially accessible (DA) cCREs between fetal and adult satellite cells (Figure 4F). The DA cCREs in fetal satellite cells were associated with biological processes such as DNA replication-dependent nucleosome assembly and triglyceride biosynthesis, and similarly to fetal skeletal myocytes were also enriched for the motifs of the MRFs Myf5 and MyoG. By contrast, adult satellite cell DA cCREs showed unexpected associations with biological processes related to regulation of hemopoiesis and immune responses, and were enriched for the binding sites of AP-1 complex members Atf3, Fos, and Fra1 (Figure 4F-G). Fetal satellite cells contained DA cCREs at genes including *MYOG* as well as *CCND2* and *RGCC*, which encode proteins involved in the regulation of myogenesis and cell cycle progression respectively (Figure 4H). Adult satellite cells, in following with ontology results related to immune system processes, contained DA cCREs located at loci encoding genes involved in inflammatory responses such *TLR4*, as well as *BMP4*, a transforming growth factor-β superfamily member with roles in embryonic development (Wang et al., 2014) that inhibits myogenic differentiation in murine muscle-derived stem cells (Wright et al., 2002). We also detected adult satellite cell DA cCREs at the locus encoding *CEBPB,* a regulator of myeloid gene expression (Huber et al., 2012) whose deficiency results in impaired muscle fiber regeneration (Marchildon et al., 2016; Ruffell et al., 2009) and whose expression in levels in peripheral blood samples correlate with muscle strength in human adults (Harries et al., 2012). Taken together, these findings reveal the regulatory elements that may underlie the proliferative capacity and quiescent nature of fetal and adult satellite cells respectively, and emphasize the value of this dataset alongside emerging human cell atlases collected at different timepoints along the lifespan for determining life stage-specific gene regulatory programs at cell type resolution.

### Chromatin features of fibroblasts in different tissue environments

Fibroblasts are the most common cells in connective tissues, and they play a critical role in orchestrating the development and morphogenesis of tissues and organs. It has become increasingly recognized that fibroblasts at different locations in the human body display distinct functions and morphologies (Chang et al., 2002; Muhl et al., 2020). However, the chromatin accessibility landscape in different fibroblast subtypes remains poorly understood. This sci-ATAC-seq dataset spanning human tissue types afforded us the opportunity to examine differences in chromatin accessibility between cellular subtypes distributed across organ systems. For example, our clustering analysis revealed six subtypes of tissue-resident fibroblasts comprised of nuclei from different tissue environments (Figure 5A). While all of these subtypes showed comparable chromatin accessibility at a set of core fibroblast cCREs, each also showed subtype-specific chromatin accessibility patterns, which were enriched for ontology terms that suggested potential subtype-specific functions (Figure 5A-B). For example, Fib.5, the fibroblast subtype derived in large proportion from sigmoid colon tissue (Figure 5A, Table S4), was enriched for biological processes related to gastrointestinal smooth muscle contraction. Fib.6, the fibroblast subtype derived mostly from hepatic and adrenal tissue – two highly-vascularized organ systems in the body, was enriched for biological processes related to positive regulation of angiogenesis (Figure 5B).

**Figure 5.**
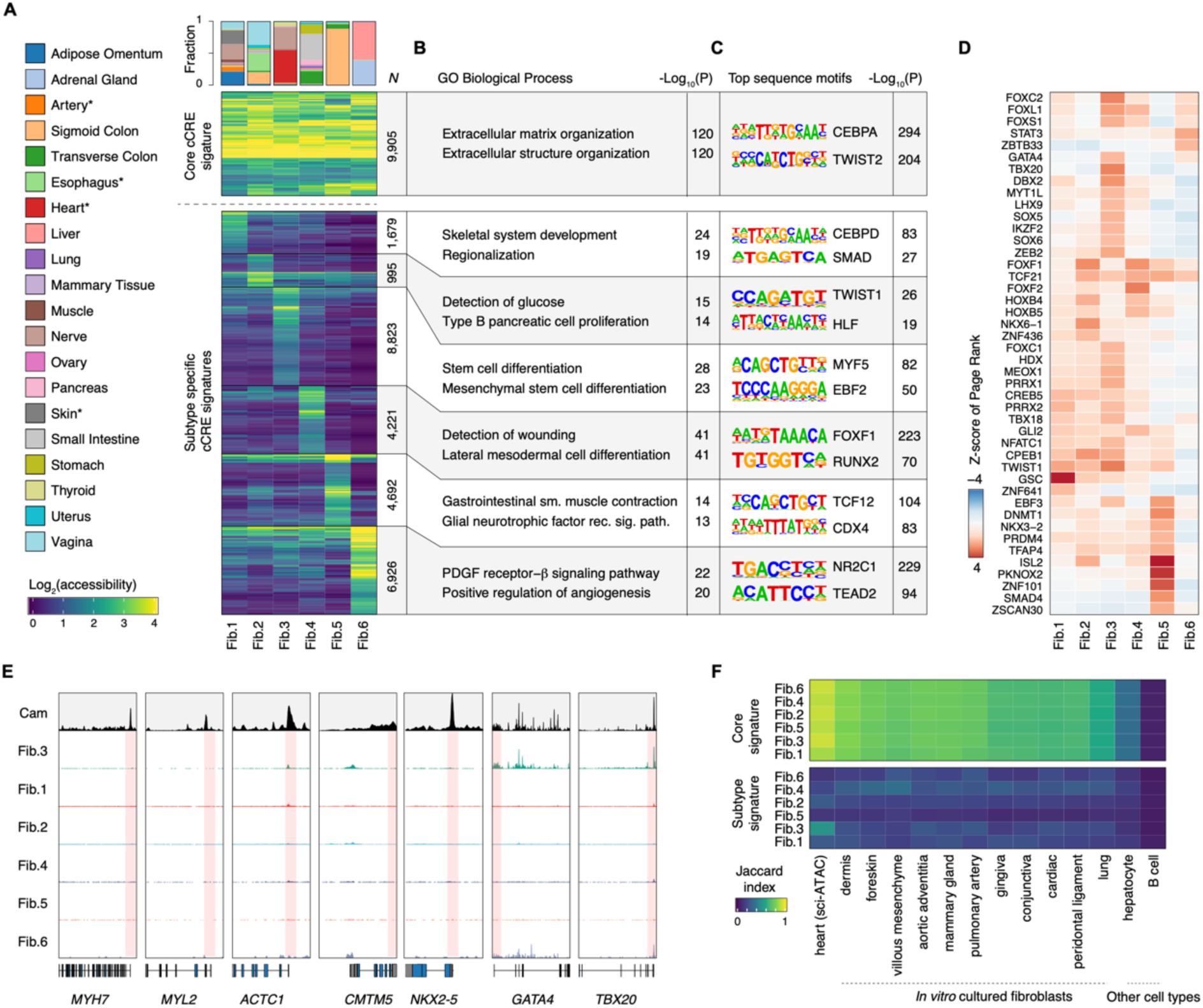
Chromatin features of fibroblasts in different tissue environments. **A)** Heatmap representation of core fibroblast cCREs and fibroblast subtype-specific elements. Color represents log_2_(accessibility). Bar plot on the top indicates tissues of origin by percentage for each fibroblast subtype. **B)** Top GREAT ontology enrichments (McLean et al., 2010) for core fibroblast and fibroblast subtype-specific cCREs. **C)** *De novo* sequence motifs and their matched known TF motifs identified by HOMER (Heinz et al., 2010). **D)** Similarity indices between (top) core fibroblast cCREs and (bottom) subtype-specific cCREs with *in vivo* cardiac fibroblasts from sci-ATAC-seq (Hocker et al., 2020), *in vitro* cultured fibroblast DNase-seq datasets, and non-fibroblast DNase-seq datasets. **E)** Heatmap representation showing key TFs (row) in each fibroblast subtype (column) revealed using transcription regulatory network analysis. Color represents standardized PageRank scores. **F)** Genome browser tracks for cardiomyocytes (Cam) and fibroblast subtypes (Fib.1-Fib.6) from sci-ATAC-seq at several cardiomyocyte marker genes.

We next examined TF motif enrichment within core and subtype-specific fibroblast cCREs. Core fibroblast cCREs were enriched for motifs of the bZIP family TF CEBPA and the bHLH family TF TWIST2 (Figure 5C). On the other hand, subtype-specific cCREs showed strong enrichments for diverse TF motifs. Encouraged by these findings, we further performed transcriptional network analysis using the PageRank algorithm (Zhang et al., 2019) to identify candidate driver TFs in each fibroblast subtype. For example, Fib.1, the fibroblast subtype derived broadly from skin, adipose, artery, skeletal muscle, and tibial nerve tissues, was enriched for the homeobox family TF GSC which is a conserved regulator of gastrulation and organogenesis in many species (Blum et al., 1992; Izpisúa-Belmonte et al., 1993; Niehrs et al., 1993) (Figure 5D). In humans, mutations in the gene encoding GSC can lead to a syndrome of short stature, auditory canal atresia, mandibular hypoplasia, and skeletal system abnormalities (Parry et al., 2013). Interestingly, Fib.3, the fibroblast subtype derived predominantly from cardiac tissue, was enriched for TFs GATA4 and TBX20 which regulate cardiac organogenesis and adult cardiomyocyte function (Perrino and Rockman, 2006; Shen et al., 2011; Singh et al., 2005). Fib.3 also showed strong accessibility at the genes encoding these TFs, but did not show accessibility at other cardiomyocyte marker genes (Figure 5E). Together, these findings are in line with recent characterizations of unexpected cardiogenic gene programs in cardiac fibroblasts (Furtado et al., 2014). We finally compared subtype-specific cCREs with chromatin profiles from *in vitro* cultured fibroblast biosamples and cardiac fibroblasts from sci-ATAC-seq (Hocker et al., 2020). While all fibroblast subtypes from the current study showed similarity to *in vitro* fibroblasts based on core fibroblast cCRE signatures, only the fibroblast subtype Fib.3 matched previously reported cardiac fibroblasts based on subtype-specific fibroblast cCRE signatures (Figure 5F), suggesting that fibroblast subtype-specific signatures are environment dependent and may be lost during *in vitro* culturing. Overall, these findings reveal a core regulatory program for adult tissue resident fibroblasts distributed across human organ systems, as well as the chromatin features and TFs that may regulate more specialized roles of tissue-resident fibroblast subtypes.

### Association of human cell types with risk variants for complex traits and diseases

Genetic variants associated with complex diseases and traits from GWAS predominantly reside in non-coding regions of the genome (Claussnitzer et al., 2020) and are enriched in cCREs in a tissue and cell type-specific fashion (Corces et al., 2020; Cusanovich et al., 2018; Domcke et al., 2020; Hocker et al., 2020; Maurano et al., 2012; Song et al., 2020; Song et al., 2019). To examine the genome-wide enrichment of disease and trait associated variants within cCREs annotated in each of the 54 human cell types characterized in the current study, we performed cell type-stratified linkage disequilibrium score regression (LDSC) analysis using GWAS summary statistics for 56 phenotypes including diseases and non-disease traits (Figure 6A-B, Table S9, See Methods). This analysis revealed a total of 163 significant associations between 38 cell types and 40 complex phenotypes (Figure 6A-B). These enrichments revealed expected cell type-disease relationships - for example, multiple sclerosis variants were strongly enriched in cCREs detected in B cells and T cells (Consortium, 2019) (False Discovery Rate (FDR) < 0.001), type 2 diabetes variants were strongly enriched in neuroendocrine cell cCREs, likely because of contributions from pancreatic beta cells (Figure S3) (Chiou et al., 2019) (FDR < 0.001), and Alzheimer’s disease variants were enriched in macrophage cCREs (FDR < 0.05) in line with their reported strong enrichment in microglial populations (Nott et al., 2019). Notably however, our analysis also revealed disease-cell type relationships for *in vivo* adult human cell types not presently annotated by bulk DNase-seq or ATAC-seq data. These included a strong enrichment of coronary artery disease variants in vascular smooth muscle cCREs (FDR < 0.01), a strong enrichment of HDL cholesterol level-associated variants in adipocyte cCREs (FDR < 0.01), and a nominal enrichment of ulcerative colitis variants in gastrointestinal goblet cell cCREs (P < 0.05) in addition to T lymphocyte cCREs (FDR < 0.01). Further, we detected differences in the enrichment of disease and trait variants in subtypes of tissue resident fibroblasts. While all fibroblast populations were enriched for variants associated with standing height to an equivalent degree (FDR < 0.001), only Fib.3, the fibroblast subtype derived mostly from heart atrial appendage and left ventricle, showed a significant enrichment for coronary artery disease variants (FDR < 0.05). Similarly, all three fibroblast subtypes with major contributions from gastrointestinal tissues including the esophagus (Fib.2), stomach and lower gastrointestinal tract (Fib.4), and sigmoid colon (Fib.5) were strongly enriched for diverticular disease-associated variants, whereas those derived mostly from cardiac tissue (Fib.3) and liver/adrenal tissue (Fib.6) were not.

**Figure 6.**
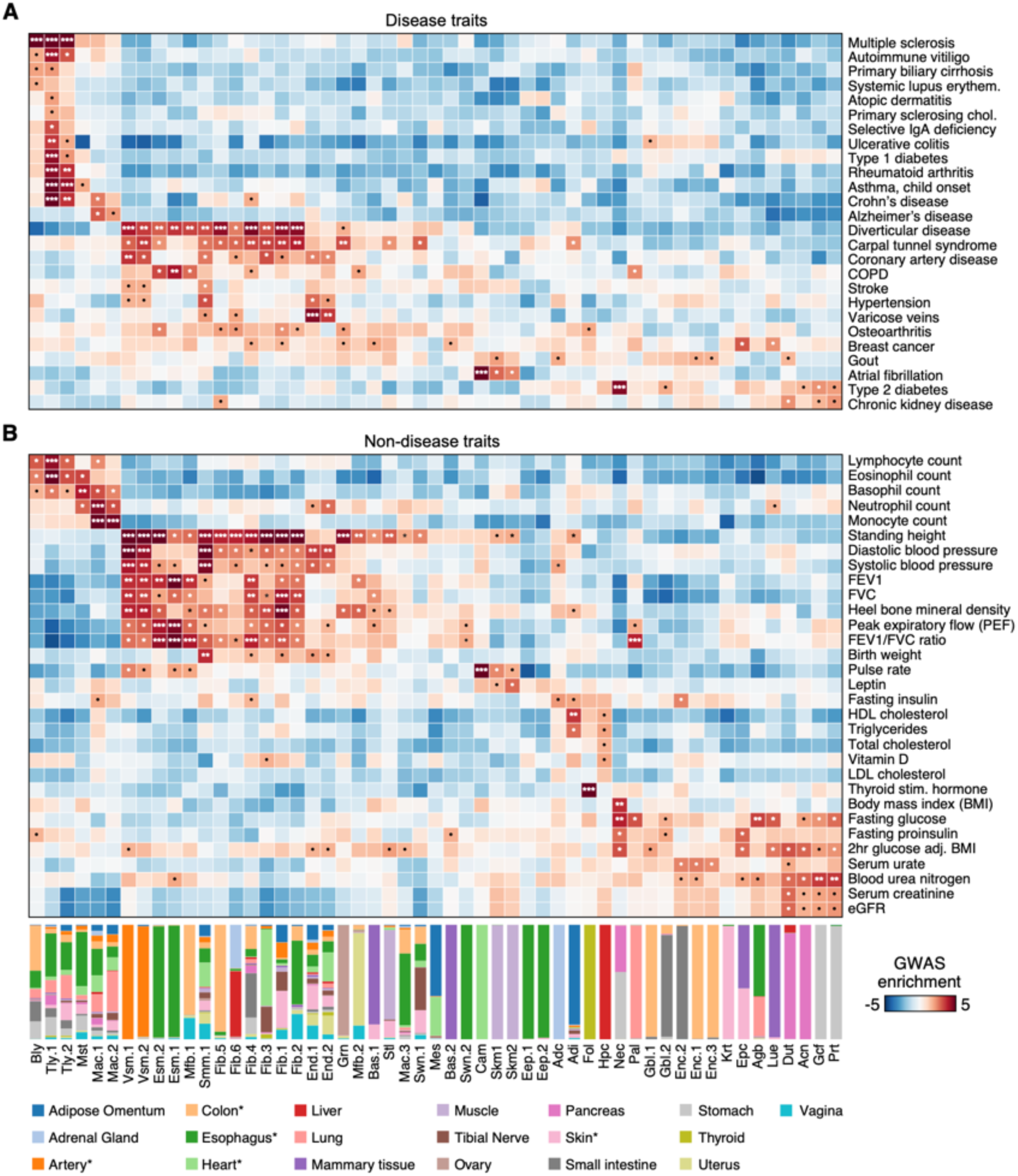
Association of human cell types with risk variants for complex traits and diseases. Heatmap showing enrichment of risk variants associated with disease (**A**) and non-disease traits (**B**) from genome wide association studies in human cell type-resolved cCREs. Cell type-stratified linkage disequilibrium score regression (LDSC) analysis was performed using GWAS summary statistics for 56 phenotypes. Total cCREs identified independently from each cell type were used as input for analysis. Z-scores for enrichment are displayed and were used to compute one-sided p-values for enrichments. P-values were corrected using the Benjamini Hochberg procedure for multiple tests (*: FDR < 0.1; **: FDR < 0.01; ***: FDR < 0.001; •: nominal p-value < 0.05). Bar plot on the bottom shows the tissue contributions for each cell cluster. * indicates categories representing multiple samples that originated from similar tissues.

### Systematic interpretation of molecular functions for non-coding risk variants

Many non-coding disease-associated genetic variants are hypothesized to alter the expression of disease-associated genes by disrupting TF binding to *cis-*regulatory elements. However, without comprehensive annotations of cCREs at cell type resolution across the human body, the molecular functions of these variants have proven challenging to interpret (Claussnitzer et al., 2020). We sought to apply our atlas of cCREs in adult human cell types to systematically interpret molecular mechanisms for genetic variants associated with complex traits and diseases.

First, we determined the probability that variants from 48 GWAS were causal for disease or trait association (Posterior probability of association, PPA) using Bayesian fine-mapping (Wakefield, 2009). We defined likely causal variants as variants with a PPA > 0.1, and found that they were more likely to reside within cCREs than the rest of the variants (Figure S11A). Overall, we detected 2,730 likely causal variants residing within cCREs mapped in various human cell types (Figure 7A-B, Table S10). Second, we analyzed previously published promoter capture HiC data in similar tissues (Jung et al., 2019) and linked our cCREs to target genes via the Activity-by-Contact (ABC) model (Fulco et al., 2019) (See Methods). This analysis revealed 3,926,564 unique distal cCRE-to-gene linkages across our 54 cell types, with a median of 760,954 total linkages and 15,680 cell type-specific linkages per cell type (Figure S11B-C; Supplementary files with distal cCRE to gene linkages downloadable from http://catlas.org/humantissue). Of the 2,730 cCREs containing likely causal variants, we linked 1,843 to putative target genes (Figure 7A). Third, we applied our recently developed deltaSVM models for 94 TFs (Yan et al., 2021) to identify the variants potentially disrupting binding by these regulators. This analysis found 460 TF binding sites that could be significantly altered by the likely causal variants (Figure 7A). The intersection of these lists prioritized 302 likely causal GWAS variants that 1) resided within a human cell type cCRE, 2) significantly altered TF binding 3) and were linked to one or more target genes (Figure 7A-B, Table S10).

**Figure 7.**
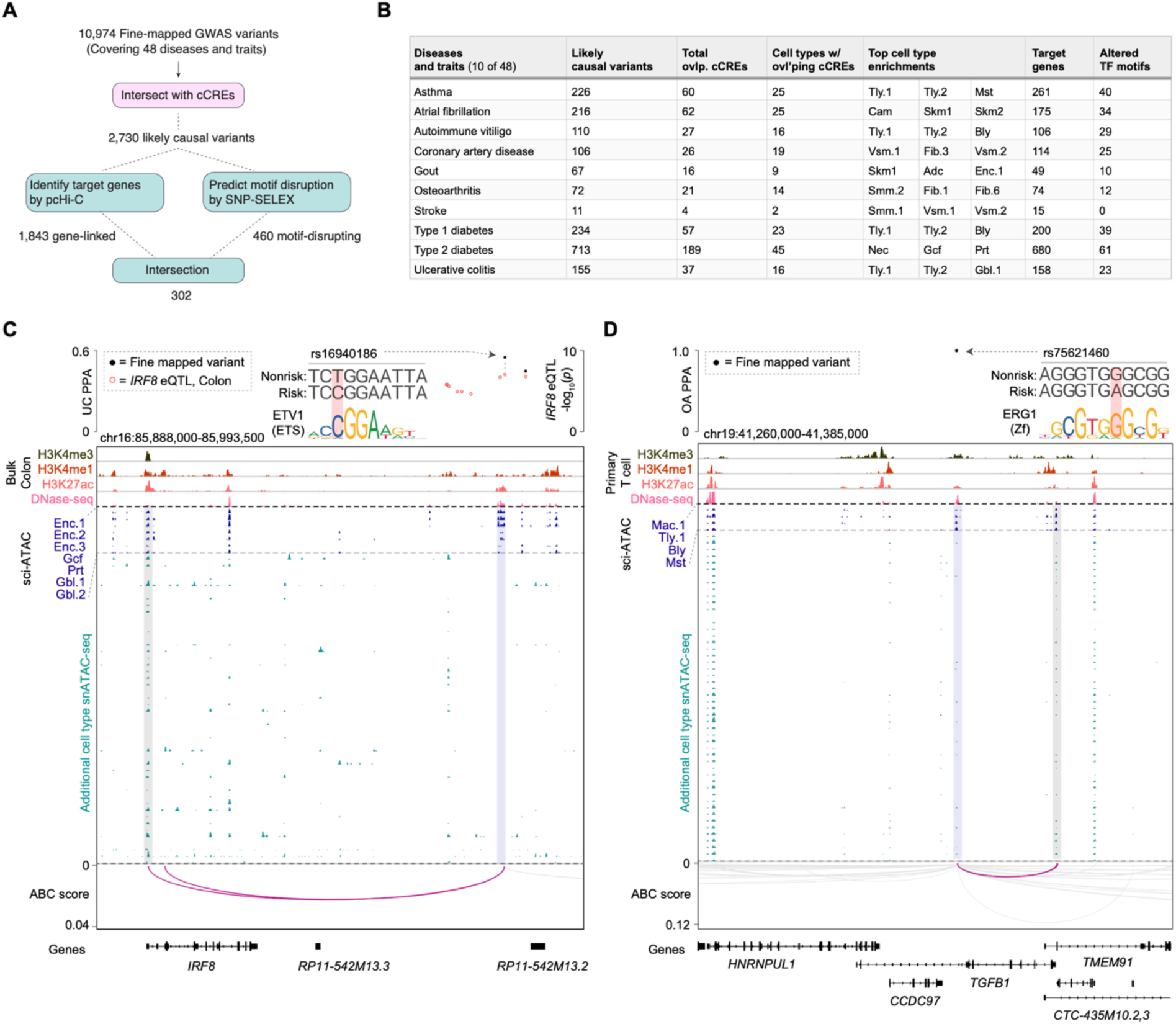
Systematic interpretation of molecular functions for non-coding risk variants. **A)** Schematic illustrating the workflow for annotating fine-mapped non-coding risk variants. We started with 10,974 likely causal fine-mapped variants (with a posterior probability of association – PPA – greater than 0.1) spanning 48 diseases or complex traits. 2,730 likely causal variants were found to overlap with human cell type cCREs defined in the present study. For each of these variants, we searched for target genes using promoter capture HiC data and identified disrupted TF motifs using 94 deltaSVM models trained using recent SNP-SELEX experiments (Yan et al., 2021). Finally, 302 likely causal variants were annotated with a full complement of information (overlapping cell type cCRE, putative target gene, and altered TF motif). **B)** Table showing for 10 examples out of 48 total fine-mapped diseases and traits: number of likely causal variants (PPA > 0.1), number of cCREs overlapping likely causal variants, number of cell types in which overlapping cCREs are accessible, top cell types variants are enriched in based on LD score regression (Bulik-Sullivan et al., 2015), number of predicted target genes for likely causal variants, and significantly altered motifs predicted by deltaSVM model trained using SNP-SELEX data. Comprehensive data are provided in Table S10. **C,D)** Fine mapping and molecular characterization of an ulcerative colitis (UC) risk variant (**C**) in a gastrointestinal (GI) epithelial cell cCRE (Enc = enterocyte, Gcf = gastric chief cell, Prt = parietal cell, Gbl = goblet cell) and an osteoarthritis variant (**D**) in an immune cell cCRE (Mac = macrophage, Tly = T lymphocyte, Bly = B lymphocyte, Mst = Mast cell). Genome browser tracks (GRCh38) display histone modification ChIP-seq and DNase-seq from public human transverse colon datasets (**C**) and human primary T cell datasets (**D**) from ENCODE (see Methods) as well as chromatin accessibility profiles for human cell types from sci-ATAC-seq. Chromatin interaction tracks show linkages between the variant-containing cCREs and genes from promoter capture HiC data via Activity-by-Contact (ABC) (Fulco et al., 2019) analysis. All linkages shown have an ABC score > 0.02. PPA: Posterior probability of association.

For example, one likely causal risk variant for ulcerative colitis (rs16940186) resided within an intergenic cCRE restricted to epithelial cells of the gastrointestinal tract including enterocytes, gastric parietal and chief cells, and goblet cells (Figure 7C). The cCRE containing rs16940186 was predicted to contact the transcription start site of *IRF8* (ABC score > 0.02), which encodes a TF involved in the regulation of immune cell maturation (Salem et al., 2020) and regulation of innate immunity in gastric epithelial cells (Yan et al., 2016). The rs16940186 risk allele is an eQTL associated with increased *IRF8* expression in human colon tissue and, consistent with these findings, SNP-SELEX motif disruption analysis predicted this risk allele to create a binding site for the ETS family of activating TFs (Figure 7C), which are expressed in intestinal epithelia and have been suggested to regulate intestinal epithelial maturation (Jedlicka et al., 2009). One other prioritized likely causal risk variant for osteoarthritis (rs75621460) resided within a cCRE that was primarily accessible in immune cell types, was predicted to target the immunosuppressive cytokine gene *TGFB1,* and disrupted a binding site for the zinc-finger TF ERG1 (Figure 7D).

## DISCUSSION

Detailed knowledge of the regulatory programs that govern gene expression in the human body has key implications for understanding human development and disease pathogenesis. Here, we used a single cell ATAC-seq method to profile chromatin accessibility in 472,373 cells across 25 adult human tissues representing a wide range of human organ systems, and to produce a cell-type resolved human cCRE atlas. The resulting maps bridge a key gap in the annotation of candidate regulatory elements in the human genome by providing state of activities of each element across 54 major cell classes. We used this atlas to reveal *cis-*regulatory programs and transcriptional regulators of adult human cell types, and characterized regulatory programs that may govern the tissue and subtype-specific functions of widely distributed cell types such as fibroblasts. We further incorporated this dataset alongside single cell chromatin accessibility data from human fetal tissues (Domcke et al., 2020), to reveal the regulatory elements that may govern life stage-specific cellular roles. The atlas of chromatin accessibility reported here is thus highly complementary to emerging atlases of chromatin accessibility in human fetal tissues (Domcke et al., 2020) and in individual human organ systems (Chiou et al., 2019; Corces et al., 2020; Hocker et al., 2020; Wang et al., 2020). Integration of these datasets along with future human single cell datasets of increasing scale, breadth, and depth will enable a comprehensive understanding of gene regulatory features of human cell types throughout the lifespan.

While genome-wide association studies (GWAS) have been broadly used to enhance our understanding of polygenic human traits and reveal clinically-relevant therapeutic targets for complex diseases, to date the discovery of new variants has far outpaced our ability to interpret their molecular functions (Claussnitzer et al., 2020). A central goal of the current study was thus to leverage novel maps of cCREs in adult human cell types to interpret the molecular functions of noncoding risk variants for complex disease. By applying our datasets alongside cutting-edge methods to prioritize likely causal variants in LD, link distal cCREs to target genes, and predict motifs altered by risk variants, we created a framework to systematically interpret noncoding risk variants and provided a resource of overlapping cCREs, associated cell types, potentially disrupted TFs, and putative gene targets for a host of fine mapped variants. For example, we highlight the likely causal ulcerative colitis-associated variant rs16940186. This risk variant may function to increase *IRF8* expression in gastrointestinal epithelial cells by creating a binding site for ETS family TFs in a GI epithelial-specific enhancer, and thereby alter the transcriptional responses of intestinal epithelial cells to inflammatory cytokines. Pending functional validation experiments, our results suggest that targeting *IRF8* in GI epithelial cells could be a potential therapeutic target for ulcerative colitis. As future GWAS in large cohorts with detailed phenotyping, whole genome sequencing efforts, and novel association studies employing long read technologies to capture structural variants become available, we anticipate that this combined resource and framework will be of continued utility for the interpretation of molecular functions for noncoding genetic variants.

The current study is still limited in several ways: firstly, we solely profiled the adult stage in an incomplete sampling of organ systems. While we utilized tissue from anatomic sites corresponding directly to existing biosamples in large-scale databases (Carithers et al., 2015; Stranger et al., 2017), the size and diversity of adult human organ systems make it difficult to representatively sample them in their entirety with current technologies. Additionally, our assay solely profiles chromatin accessibility in dissociated nuclei, and thus misses key orthogonal molecular and spatial information. Future assays that incorporate gene expression, chromatin accessibility, DNA methylation, chromosomal conformation, TF binding, and spatial information in the same single cell will greatly enhance our understanding of gene regulation in human cell types (Zhu et al., 2020). Notwithstanding these limitations, this atlas of >750,000 cCREs in almost half a million nuclei represents the largest cellular survey of cCREs across adult human organ systems to the best of our knowledge. This resource thus lays the foundation for the analysis of gene regulatory programs across human organ systems at cell type resolution, and accelerates the interpretation of noncoding sequence variants associated with complex human diseases and phenotypes. The datasets can be accessed and explored at http://catlas.org/humantissue.

## ACKNOWLEDGEMENTS

We thank the ENCODE consortium, in particular Mike Pazin (NHGRI) and Idan Gadbank (Stanford), Kristin Ardlie (Broad Institute) and Ellen Gelfand (Broad Institute), for providing the tissue samples for the present study. We thank B. Li for bioinformatics support. We thank S. Kuan for sequencing libraries on the HiSeq4000. We thank B. Chen for valuable discussions and feedback. We thank the QB3 Macrolab at UC Berkeley for purification of the Tn5 transposase. This work was supported by the Ludwig Institute for Cancer Research (B.R.), and Foundation for the National Institutes of Health (K.J.G). J.D.H. was supported in part by a Ruth L. Kirschstein Institutional National Research Service Award T32 GM008666 from the National Institute of General Medical Sciences. Work at the Center for Epigenomics was supported in part by the UC San Diego School of Medicine.

## AUTHOR CONTRIBUTIONS

Study was conceived by: J.D.H., S.P., A.W., and B.R. Study supervision: B.R. Supervision of data generation: S.P., A.W. and B.R. Contribution to data generation: J.D.H., X.H., M.M. Contribution to data analysis: K.Z., J.D.H., J.C., O.P. Y.E.L., Y.Q. Contribution to web portal: Y.E.L., K.Z. Contribution to data interpretation: K.Z., J.D.H., S.P., A.W., K.J.G. Contribution to writing the manuscript: K.Z., J.D.H., B.R. All authors edited and approved the manuscript.

## SUPPLEMENTAL FIGURES

**Supplemental Figure 1.**
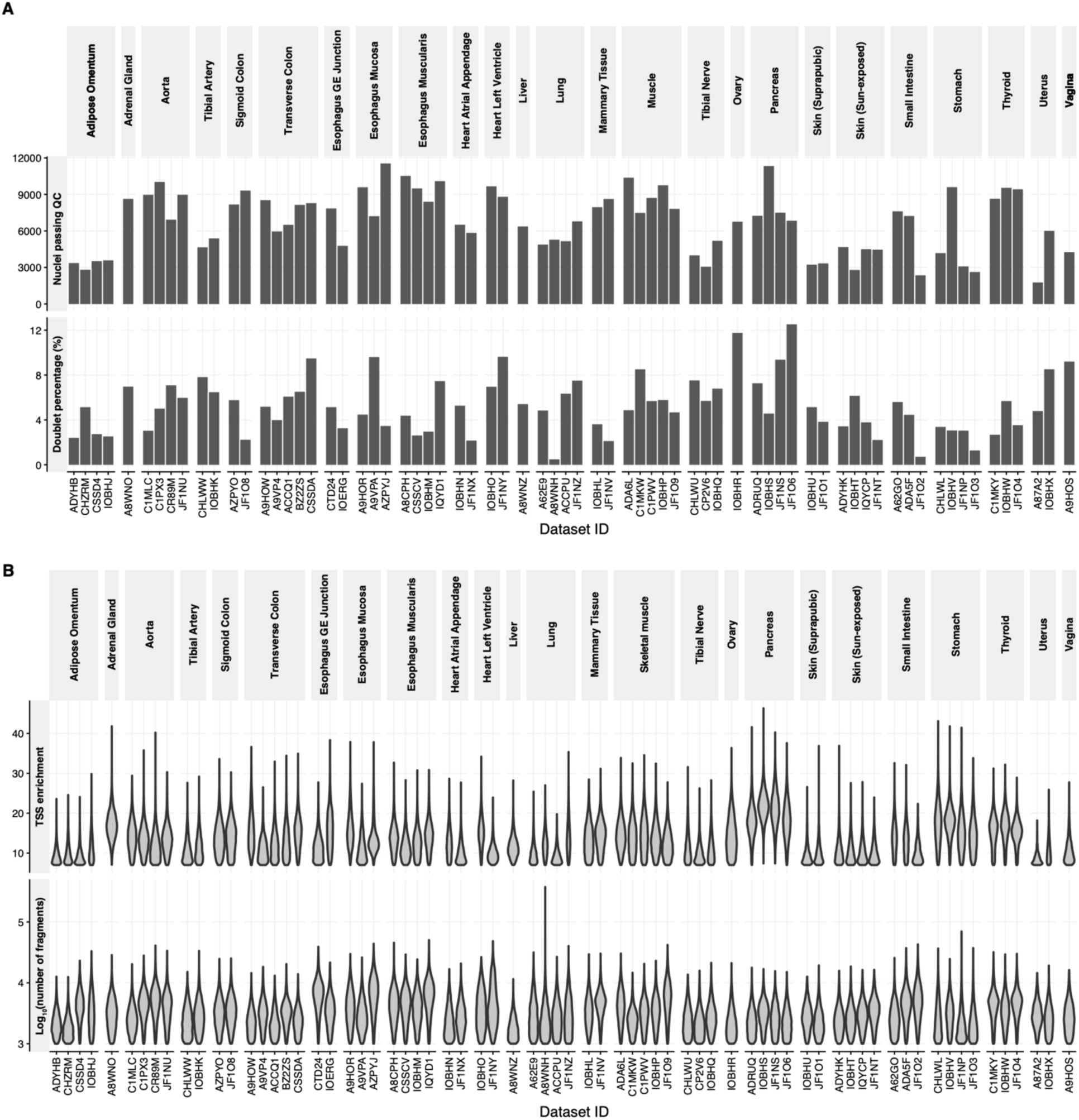
Quality control for sci-ATAC-seq datasets. **A)** Upper bar plot shows the number of nuclei that passed quality control in each experiment. Nuclei were first filtered by stringent quality control criteria (TSS enrichment greater than 7 and number of mapped fragments greater than 1000 per nucleus) and then subjected to doublet removal. Lower bar plot bottom shows the percentage of doublets detected in each dataset. **B)** Upper violin plot shows the distribution of TSS enrichments for nuclei that passed quality control in each experiment. Lower violin plot shows the distribution of number of fragments for nuclei that passed quality control in each dataset.

**Supplemental Figure 2.**
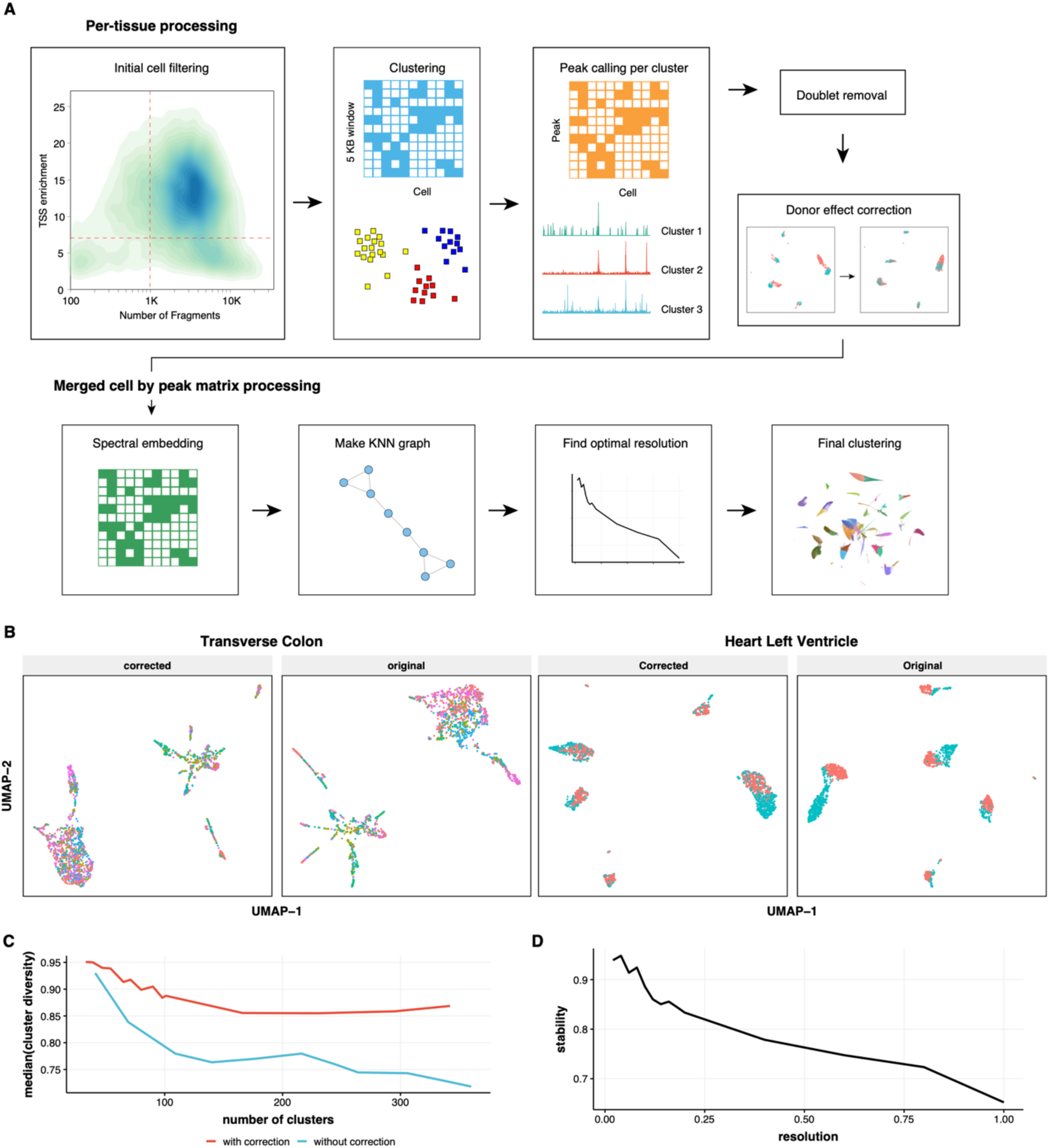
Computational framework for analyzing sci-ATAC-seq data. **A)** Schematic illustrating the workflow of the analysis pipeline. **B)** Scatter plots showing the UMAP embedding of nuclei before and after batch correction. Dots with the same color are coming from the same donor or batch. **C)** Line plot showing the median of cluster diversity as a function of number of identified clusters in the dataset stratified by batch correction operation. To compute the cluster diversity, we first grouped the cells based on their tissue of origin and then based on the experimental batch. We counted the cells for each combination and normalized by the total number of cells of the corresponding sample. For each tissue, normalized entropy was computed across batches. The average entropy across all tissues in the cluster were taken as the cluster diversity. **D)** Line plot showing the stability of clustering results as a function of resolution parameter in the Leiden algorithm. To compute the stability under a particular resolution, five perturbations were conducted on the kNN graph. During each perturbation 2% of the edges were randomly selected and subject to removal. The clustering was performed on the perturbed graph and the average Adjusted Rand Index (ARI) between different runs were taken as the stability.

**Supplemental Figure 3.**
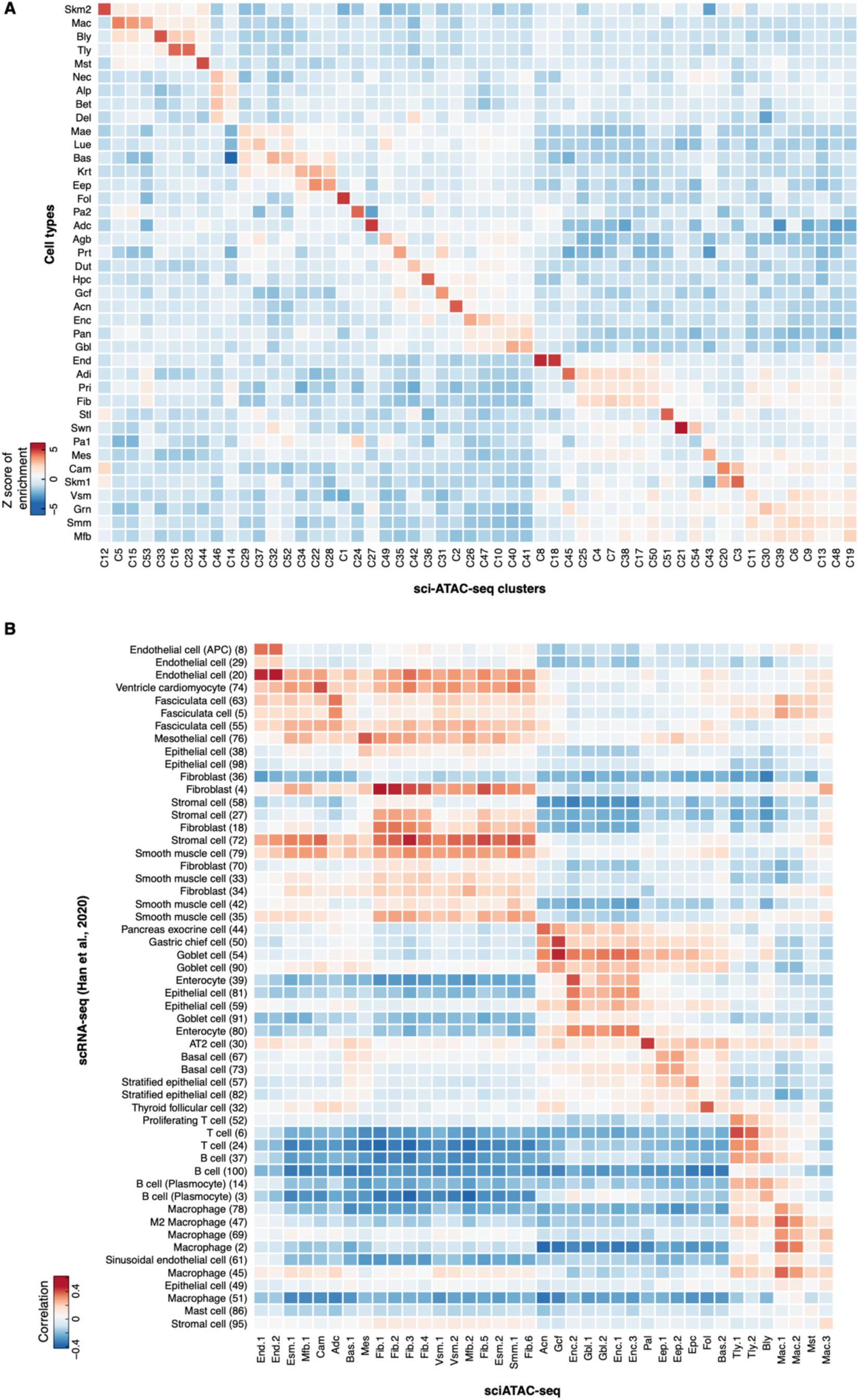
Evidence supporting the annotation of 54 cell clusters. **A)** Heatmap representation showing the marker gene enrichment of cell types. The marker genes were downloaded from the PanglaoDB (Franzén et al., 2019). **B)** Heatmap representation showing the pairwise similarity between 39 sci-ATAC-seq cell types (column) and corresponding scRNA-seq cell types (row). Color represents the Pearson correlation coefficient of expression level of 500 most variable genes. Promoter accessibility was used to estimate the gene expression level in sci-ATAC-seq.

**Supplemental Figure 4.**
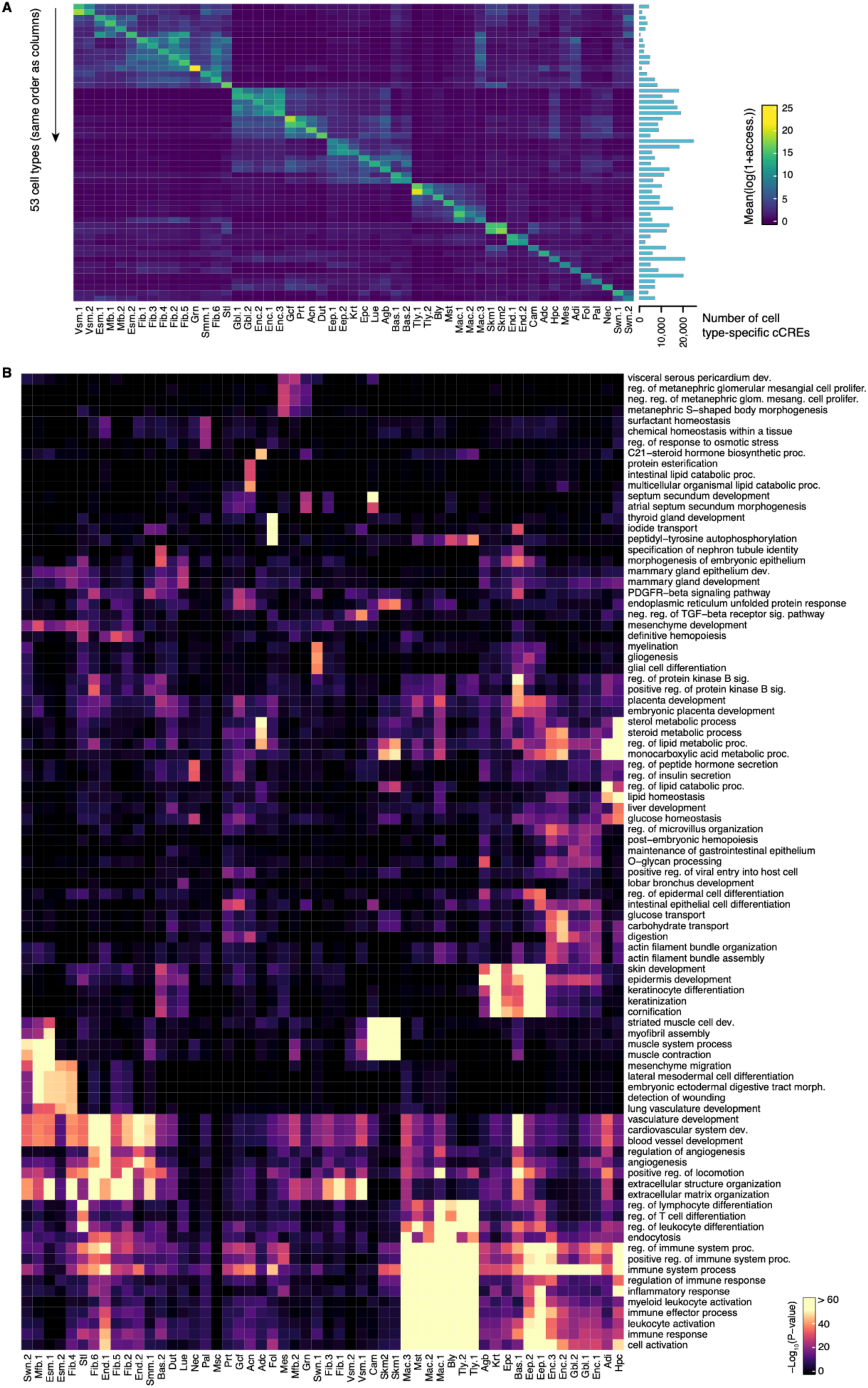
Characterization of cell-type-restricted cCREs in 53 out of 54 sci-ATAC-seq cell types. **A)** Chromatin accessibility at cell type-specific cCREs. Color represents the average log_2_(accessibility) of the cell-type-restricted cCREs in a particular cell type. Each row represents the aggregated profile of cell-type-restricted cCREs. Bar plot on the right shows the number of cell type-specific cCREs for each cell type. **B)** Heatmap representation showing the gene ontology term (column) enrichment for each set of cell-type-restricted cCREs (row). The enrichment analysis was performed using GREAT (McLean et al., 2010) under default settings. Color represents the negative logarithm of P-value of enrichment.

**Supplemental Figure 5.**
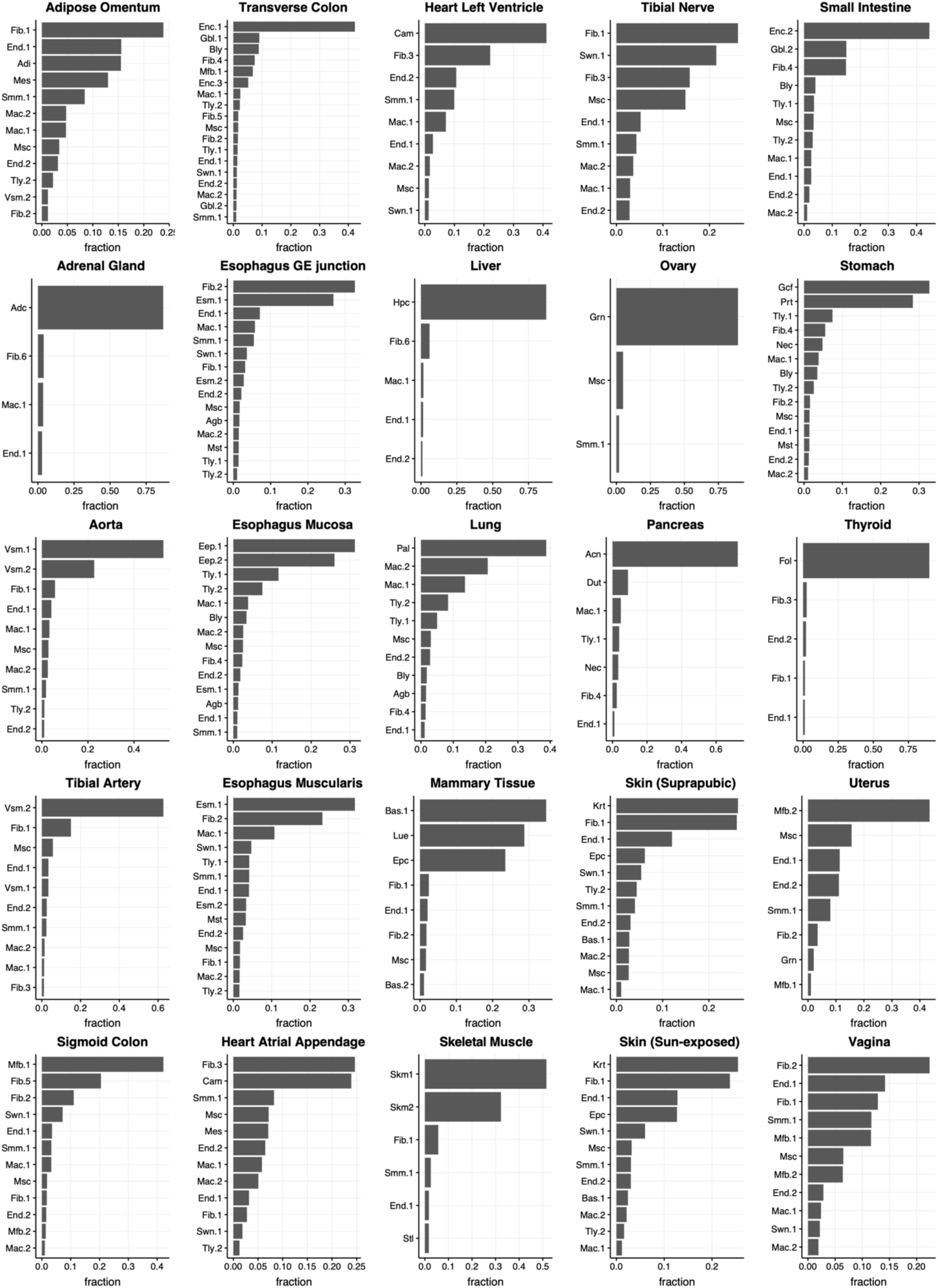
Bar plots showing cell-type composition for 25 tissue types.

**Supplemental Figure 6.**
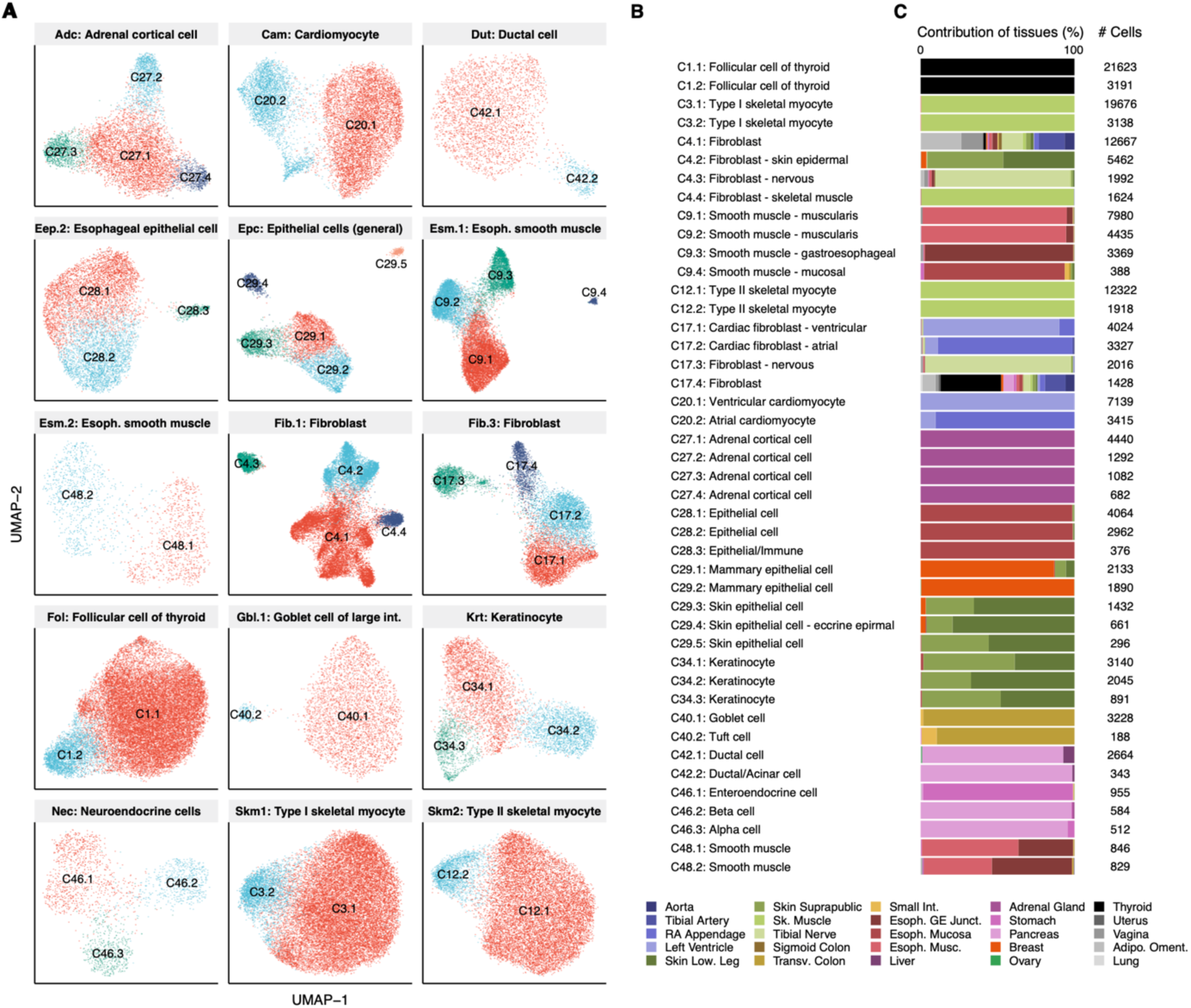
Focused clustering analysis reveals heterogeneity in primary cell clusters. **A)** UMAP embedding of cells from 15 primary cell clusters that contain more than one subcluster during focused clustering analysis. **B)** Cell type annotation of 44 subclusters based on chromatin accessibility at marker genes. **C)** Bar chart showing relative contributions of tissues to 44 subclusters.

**Supplemental Figure 7.**
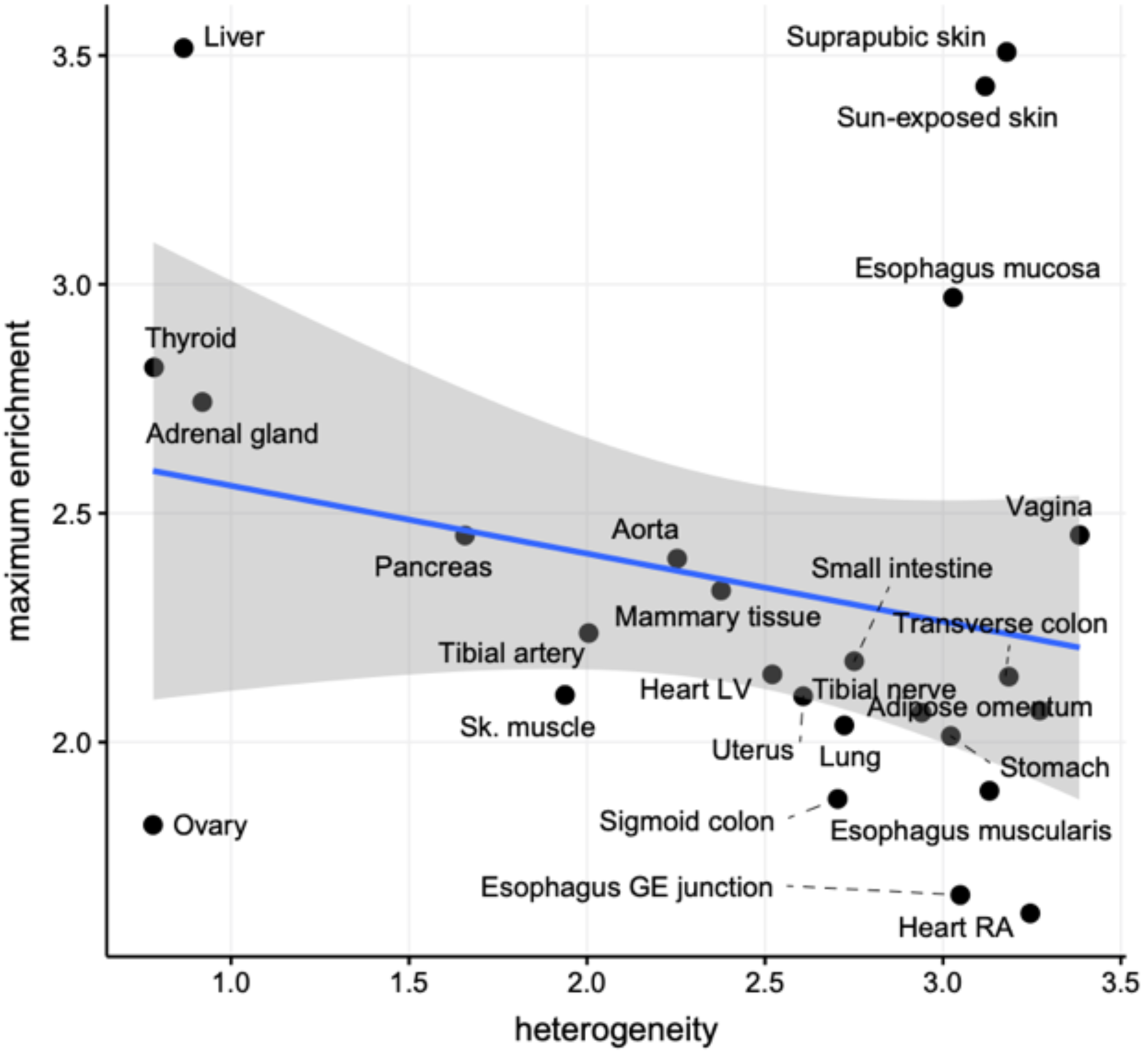
Scatter plot showing the maximum chromatin accessibility enrichment of GTEx tissue eQTLs as a function of cellular heterogeneity. The chromatin accessibility enrichment of GTEx tissue eQTLs in each tissue was computed as described in Method, and the maximum value across the 25 tissue types was used for the plot.

**Supplemental Figure 8.**
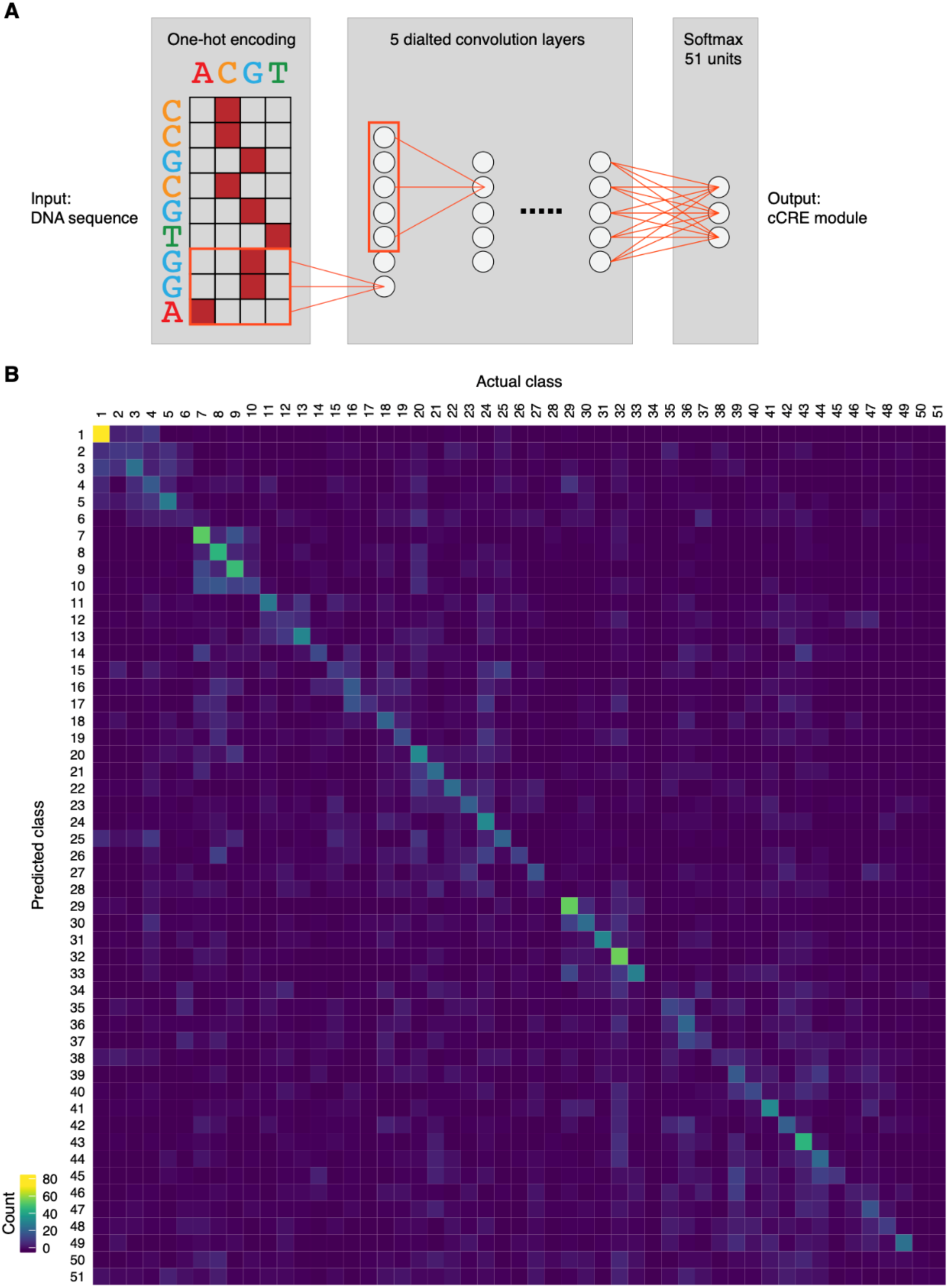
Convolutional neural network identifies sequence determinants of regulatory modules. **A)** Schematic illustrating the architecture of a 51-class neural network consisting of 5 dilated convolutional neural network layers. **B)** Heatmap representation of the confusion matrix. Each row of the matrix represents the instances in a predicted class while each column represents the instances in an actual class. Color represents the number of CREs.

**Supplemental Figure 9.**
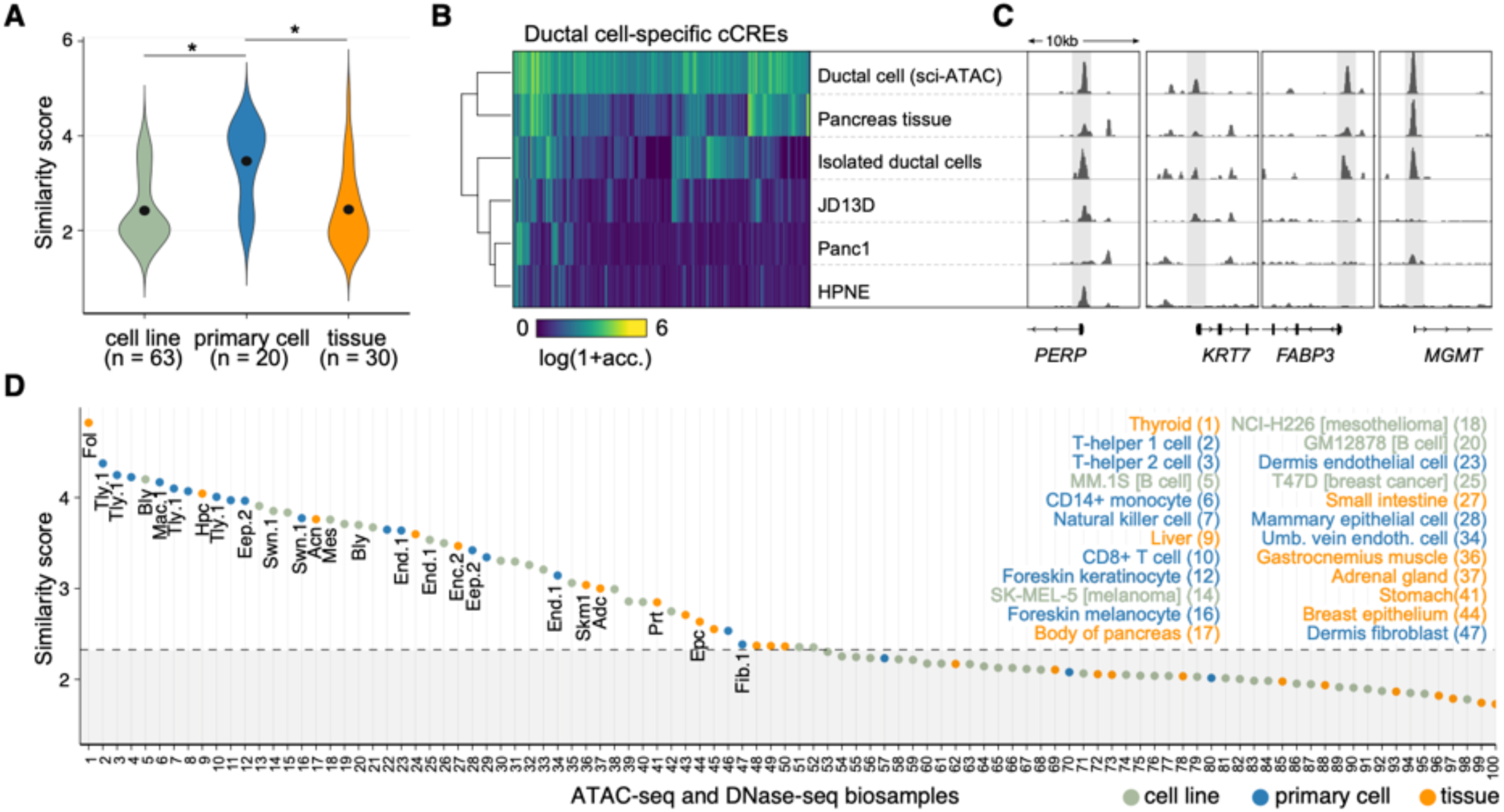
Comparison of open chromatin landscapes in adult human cell types with previous DNase-seq data obtained from bulk biosamples. **A)** Distribution of similarity scores for 113 bulk DNase-seq samples stratified by sample classification. Similarity score is defined as the maximum of the standardized correlation scores of a bulk DNase-seq sample with 54 adult human cell types from sci-ATAC-seq. * indicates P value < 0.01. Green color denotes data from cell lines, blue color denotes data from primary cells, and orange color denotes data from bulk tissues. **B)** Heatmap representation of chromatin accessibility at ductal cell-specific cCREs identified by sci-ATAC-seq across ductal cell-related sci-ATAC-seq, primary cell, tissue, and immortal cell line biosamples. **C)** genome browser tracks showing chromatin accessibility profiles around ductal cell marker genes (*PERP* and *KRT7*) or tumor repressors (*FABP3* and *MGMT*). **D)** Top similarity scores by rank shown for 100 bulk biosamples & corresponding best match cell types. Sample classification is indicated by color.

**Supplemental Figure 10.**
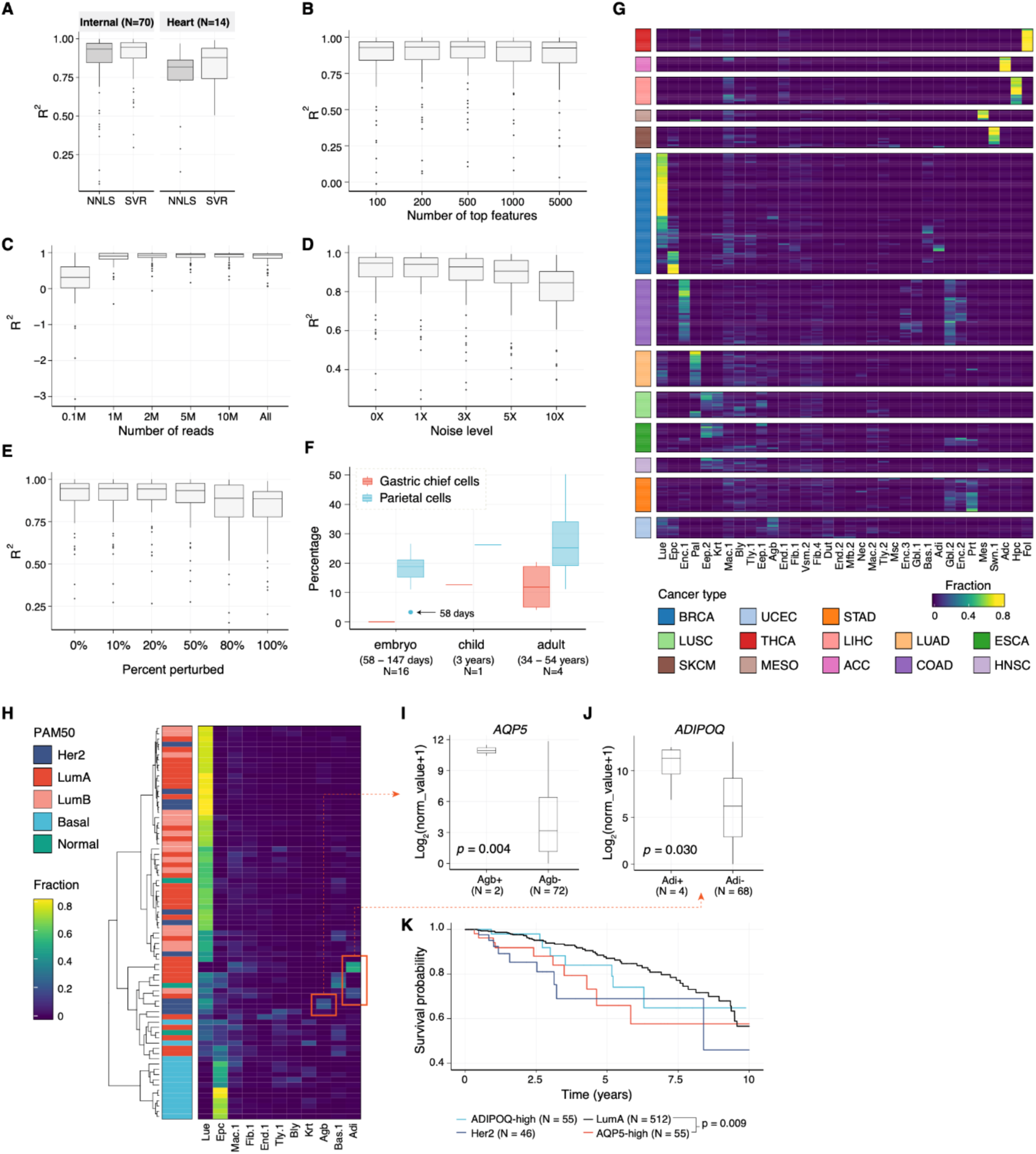
CRE cytometry reveals tissue heterogeneity of primary human cancer. **A)** Boxplot showing the performance of two deconvolution algorithms, namely non-negative least squares regression (NNLS) and support vector regression (SVR). The performance is measured by coefficient of determination (*R*^!^) between estimated cell-type composition and actual cell-type composition determined by sci-ATAC-seq experiments. In addition to the dataset generated in this study, referred to as “internal”, we performed benchmarking using independent sci-ATAC-seq datasets from 14 heart (Hocker et al., 2020). **B)** Boxplot showing the performance of NNLS, measured by coefficient of determination, under different choices of signature CREs. For example, “100” indicates selecting top 100 most specific CREs from each cell types. **C)** Boxplot showing the performance of SVR under different rates of down sampling. **D)** Boxplot showing the performance of SVR under different noise levels. For example, “1X” indicates introducing 100% more noise to the data. **E)** Boxplot showing the performance of SVR when introducing noise to a random subset of the signature CREs. The noise level here is fixed to “10X”. **F)** Boxplot showing estimated cell-type composition of 21 human stomach tissue stratified by life stage. The deconvolution was performed on bulk DNase-seq data using the SVR algorithm. **G)** Heatmap representation of cell-type composition of 275 cancer samples from TCGA. Color represents cell-type fraction. Color bars to the left depict the cancer type (BRCA = Breast invasive carcinoma, LUSC = Lung squamous cell carcinoma, SKCM = Skin cutaneous melanoma, UCEC = Uterine corpus endometrial carcinoma, THCA = Thyroid carcinoma, MESO = Mesothelioma, STAD = Stomach adenocarcinoma, LIHC = Liver hepatocellular carcinoma, ACC = Adrenocortical carcinoma, LUAD = Lung adenocarcinoma, COAD = Colon adenocarcinoma, ESCA = Esophageal carcinoma, HNSC = Head and neck squamous cell carcinoma). The deconvolution was performed on bulk ATAC-seq data using the SVR algorithm. **H)** Heatmap representation of cell-type composition of 75 breast cancer samples. Color represents cell-type fraction. The dendrogram was generated by hierarchical clustering. Published PAM50 classification scheme (Berger et al., 2018) is shown on the left. **I)** Boxplot showing the AQP5 gene expression level in breast cancer samples stratified by the presence of airway goblet cell signature. **J)** Boxplot showing the ADIPOQ gene expression level in breast cancer samples stratified by the existence of adipocyte signature. **K)** Kaplan-Meier analysis of overall survival of breast cancer sample donors in four subtype groups: LumA (N=512), AQP5 overexpressed (N=55), ADIPOQ overexpressed (N=55) and Her2 (N=46).

**Supplemental Figure 11.**
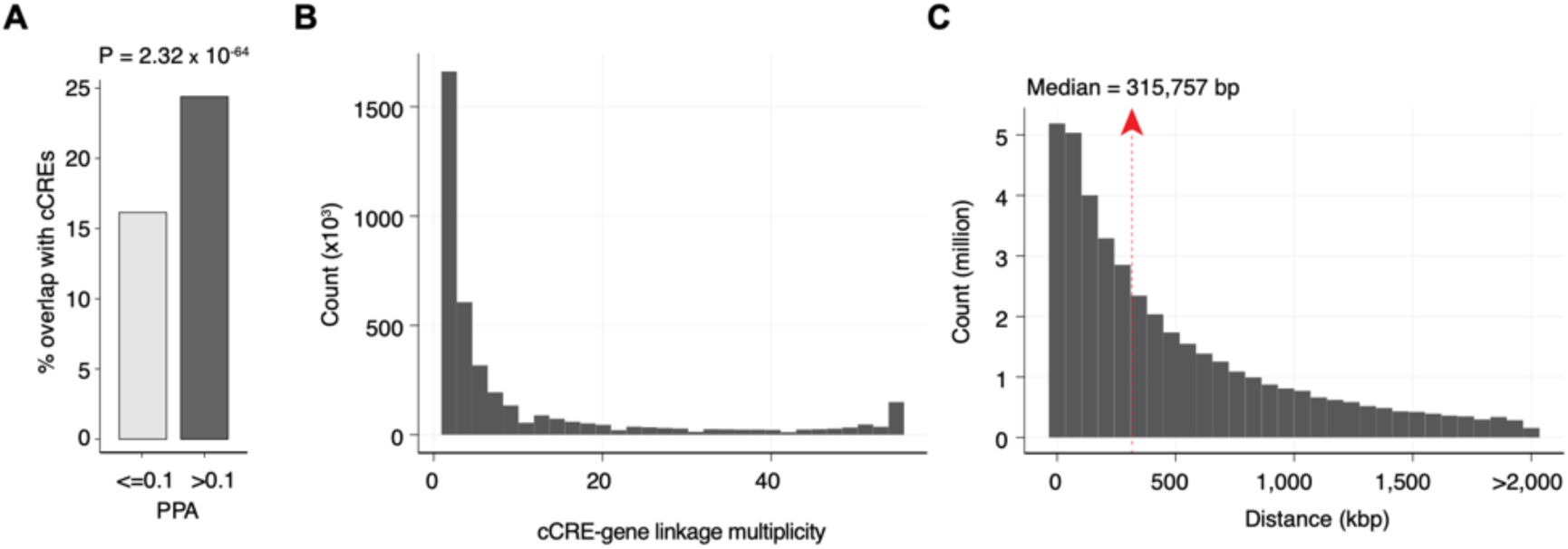
Characterization of fine mapped risk variant. **A)** Bar graph showing the percentage of likely causal (Posterior Probability of Association; PPA > 0.1) fine mapped GWAS variants from 48 traits and diseases that overlap the union set of cCREs in adult cell types in the present study. Fisher’s exact test was used to compute statistical significance. **B)** Histogram showing the multiplicities of cCRE-gene linkage (number of cell types having the linkage). **C)** Histogram showing distances in kilobase pairs (kbp) for distal cCRE-to-gene linkages from Activity by Contact (ABC) analysis (Fulco et al., 2019) (ABC score > 0.02).

## SUPPLEMENTARY TABLES

Table S1: Donor clinical characteristics and contributions to sci-ATAC-seq datasets.

Table S2: Feasibility testing results for primary human tissue types.

Table S3: Quality control data for sci-ATAC-seq datasets.

Table S4: Clustering information and quality control data for sci-ATAC-seq nuclei.

Table S5: Cell type annotations and example marker genes.

Table S6: Union set of cCREs.

Table S7: GREAT ontology results for cCRE modules.

Table S8: Similarity scores for bulk ATAC-seq and DNase-seq biosamples.

Table S9: GWAS LDSC enrichment Z-scores and P-values.

Table S10: PPAs, overlapping cCREs, corresponding cell types, motifs altered, and candidate target genes for likely causal GWAS variants.

Table S11: Oligo and primer sequences for sci-ATAC-seq.

Table S12: Primer sequences for feasibility test RT-PCR.

## METHODS

### Human Tissues

Adult human tissue samples were acquired by the ENTEx collaborative project (Stranger et al., 2017) via the GTEx collection pipeline (Carithers et al., 2015). All human donors were deceased, and informed consent was obtained via next-of-kin consent for the collection and banking of deidentified tissue samples for scientific research. Donor eligibility requirements were as described previously (Carithers et al., 2015), and excluded individuals with metastatic cancer and individuals who had received chemotherapy for cancer within the prior two years.

### Tissue feasibility testing for sci-ATAC-seq

Frozen tissue samples were sectioned on dry ice into two aliquots of equivalent mass. For nuclear isolation, one aliquot was subjected to manual pulverization via mortar and pestle while submerged in liquid nitrogen, and the other aliquot was homogenized in a gentleMACS M-tube (Miltenyi) on a gentleMACS Octo Dissociator (Miltenyi) using the “Protein_01_01” protocol in MACS buffer (5 mM CaCl2, 2 mM EDTA, 1X protease inhibitor (Roche, 05-892-970-001), 300 mM MgAc, 10 mM Tris-HCL pH 8, 0.6 mM DTT) and pelleted with a swinging bucket centrifuge (500 x g, 5 min, 4°C; 5920R, Eppendorf). Pulverized frozen tissue and pelleted nuclei from gentleMACS M-tubes were each split into two further aliquots. One aliquot from each of the two nuclear isolation conditions was then resuspended in 1 mL Nuclear Permeabilization Buffer (1X PBS, 5% Bovine Serum Albumin, 0.2% IGEPAL CA-630 (Sigma), 1 mM DTT, 1X Protease inhibitor), and the other aliquot from the same nuclear isolation condition was resuspended in 1 mL OMNI Buffer (10mM Tris-HCL (pH 7.5), 10mM NaCl, 3mM MgCl2, 0.1% Tween-20 (Sigma), 0.1% IGEPAL-CA630 (Sigma) and 0.01% Digitonin (Promega) in water), yielding a total of four nuclear isolation/nuclear permeabilization buffer conditions tested for each tissue type. Nuclei were rotated at 4 °C for 5 minutes before being pelleted again with a swinging bucket centrifuge (500 x g, 5 min, 4°C; 5920R, Eppendorf). After centrifugation, permeabilized nuclei were resuspended in 500 μL high salt tagmentation buffer (36.3 mM Tris-acetate (pH = 7.8), 72.6 mM potassium-acetate, 11 mM Mg-acetate, 17.6% DMF) and counted using a hemocytometer. Concentration was adjusted to 2,000 nuclei/9 μl, and 2,000 nuclei were dispensed 12 wells of a 96-well plate per nuclear isolation/permeabilization condition (samples were processed in batches of 4 nuclear isolation/permeabilization conditions per 2 different tissue samples). For tagmentation, 1 μL barcoded Tn5 transposomes (Table S11) were added using a BenchSmart™ 96 (Mettler Toledo), mixed five times, and incubated for 60 min at 37 °C with shaking (500 rpm).

To inhibit the Tn5 reaction, 10 µL of 40 mM EDTA (final 20mM) were added to each well with a BenchSmart™ 96 (Mettler Toledo) and the plate was incubated at 37 °C for 15 min with shaking (500 rpm). Next, 20 µL of 2x sort buffer (2 % BSA, 2 mM EDTA in PBS) were added using a BenchSmart™ 96 (Mettler Toledo). All 12 wells from each nuclear isolation/permeabilization condition were combined into a separate FACS tube, and stained with Draq7 at 1:150 dilution (Cell Signaling). For each nuclear isolation/permeabilization condition, we used a SH800 (Sony) to sort four wells containing 0 nuclei per well and four wells containing 80 nuclei per well into one 96-well plate (total of 768 wells) containing 10.5 µL EB (25 pmol primer i7, 25 pmol primer i5, 200 ng BSA (Sigma)). After addition of 1 µL 0.2% SDS using a BenchSmart™ 96 (Mettler Toledo), the 96 well plate was incubated at 55 °C for 7 min with shaking (500 rpm). 1 µL 12.5% Triton-X was added to each well to quench the SDS. Next, 12.5 µL NEBNext High-Fidelity 2× PCR Master Mix (NEB) were added to each well and samples were PCR-amplified (72 °C 5 min, 98 °C 30 s, (98 °C 10 s, 63 °C 30 s, 72°C 60 s) × 12 cycles, held at 12 °C). After PCR, all wells were assayed for DNA library concentration using the PerfeCTa NGS Quantification RT-qPCR Kit (Quanta Biosciecnces) according to manufacturer’s protocols, and subsequently returned to the thermal cycler for a second round of PCR amplification (72 °C 5 min, 98 °C 30 s, (98 °C 10 s, 63 °C 30 s, 72°C 60 s) × 4 cycles, held at 12 °C). After the second PCR amplification, for each nuclear isolation/permeabilization condition, wells containing 0 nuclei were combined and wells containing 80 nuclei were combined. The resulting DNA libraries were purified according to the MinElute PCR Purification Kit manual (Qiagen) and size selection was performed with SPRISelect reagent (Beckmann Coulter, 0.55x and 1.5x). Final libraries were quantified using a Qubit fluorimeter (Life technologies) and a nucleosomal pattern of fragment size distribution was verified using a Tapestation (High Sensitivity D1000, Agilent). We calculated a signal to noise ratio for final feasibility test libraries using LightCycler® 480 SYBR Green I Master Mix (Roche) along with custom primers for the promoter of human *GAPDH* and a heterochromatic gene desert region (Table S12). For each tissue type, the nuclear isolation/permeabilization condition that resulted in optimized nuclear yield (nuclei/mg tissue), library concentrations > 50 pM per 80 sorted nuclei, nucleosomal distribution pattern of fragments, and a log_2_(signal to noise ratio) > 3.3 was selected for combinatorial indexing-assisted single nucleus ATAC-seq (Table S2).

### Combinatorial indexing-assisted single nucleus ATAC-seq

Combinatorial indexing-assisted single nucleus ATAC-seq was performed as described previously (Preissl et al., 2018) with slight modifications (Hocker et al., 2020). Nuclei were isolated and permeabilized according to the optimized conditions from feasibility testing (Table S2). After resuspension in permeabilization buffer, nuclei were rotated at 4 °C for 5 minutes before being pelleted again with a swinging bucket centrifuge (500 x g, 5 min, 4°C; 5920R, Eppendorf). After centrifugation, permeabilized nuclei were resuspended in 500 μL high salt tagmentation buffer (36.3 mM Tris-acetate (pH = 7.8), 72.6 mM potassium-acetate, 11 mM Mg-acetate, 17.6% DMF) and counted using a hemocytometer. Concentration was adjusted to 2,000 nuclei/9 μl, and 2,000 nuclei were dispensed into each well of a 96-well plate per sample (96 tagmentation wells/sample, samples were processed in batches of 2-4 samples). For tagmentation, 1 μL barcoded Tn5 transposomes (Table S11) were added using a BenchSmart™ 96 (Mettler Toledo), mixed five times, and incubated for 60 min at 37 °C with shaking (500 rpm). To inhibit the Tn5 reaction, 10 µL of 40 mM EDTA (final 20mM) were added to each well with a BenchSmart™ 96 (Mettler Toledo) and the plate was incubated at 37 °C for 15 min with shaking (500 rpm). Next, 20 µL of 2x sort buffer (2 % BSA, 2 mM EDTA in PBS) were added using a BenchSmart™ 96 (Mettler Toledo). All wells were combined into a separate FACS tube for each sample, and stained with Draq7 at 1:150 dilution (Cell Signaling). Using a SH800 (Sony), 20 nuclei per sample were sorted per well into eight 96-well plates (total of 768 wells) containing 10.5 µL EB (25 pmol primer i7, 25 pmol primer i5, 200 ng BSA (Sigma)). Preparation of sort plates and all downstream pipetting steps were performed on a Biomek i7 Automated Workstation (Beckman Coulter). After addition of 1 µL 0.2% SDS, samples were incubated at 55 °C for 7 min with shaking (500 rpm). 1 µL 12.5% Triton-X was added to each well to quench the SDS. Next, 12.5 µL NEBNext High-Fidelity 2× PCR Master Mix (NEB) were added and samples were PCR-amplified (72 °C 5 min, 98 °C 30 s, (98 °C 10 s, 63 °C 30 s, 72°C 60 s) × 12 cycles, held at 12 °C). After PCR, all wells were combined. Libraries were purified according to the MinElute PCR Purification Kit manual (Qiagen) using a vacuum manifold (QIAvac 24 plus, Qiagen) and size selection was performed with SPRISelect reagent (Beckmann Coulter, 0.55x and 1.5x). Libraries were purified one more time with SPRISelect reagent (Beckman Coulter, 1.5x). Libraries were quantified using a Qubit fluorimeter (Life technologies) and a nucleosomal pattern of fragment size distribution was verified using a Tapestation (High Sensitivity D1000, Agilent). Libraries were sequenced on a NextSeq500 or HiSeq4000 sequencer (Illumina) using custom sequencing primers with following read lengths: 50 + 10 + 12 + 50 (Read1 + Index1 + Index2 + Read2). Primer and index sequences are listed in Table S11.

### Demultiplexing of single nucleus ATAC-seq sequencing reads

For each sequenced single nucleus ATAC-Seq library, we obtained four FASTQ files, two for paired end DNA reads and two for the combinatorial indexes for i5 and T7 (768 and 364 indices, respectively). We selected all reads with up to 2 mismatches per i5 and T7 index (Hamming distance between each pair of indices is 4) and integrated the concatenated barcode at the beginning of the read name in the demultiplexed FASTQ files. The customized scripts can be found at: https://gitlab.com/Grouumf/ATACdemultiplex/.

### Quality control metrics: TSS enrichment and unique fragments

TSS positions were obtained from the GENCODE database v31 (Frankish et al., 2019). Tn5 corrected insertions were aggregated ±2000 bp relative (TSS strand-corrected) to each unique TSS genome wide. Then this profile was normalized to the mean accessibility ± (1900 to 2000) bp from the TSS and smoothed every 11 bp. The max of the smoothed profile was taken as the TSS enrichment. We then filtered out all single cells that had fewer than 1,000 unique fragments and/or a TSS enrichment of less than 7 for all data sets.

### Overall clustering strategy

We utilized two rounds of clustering analysis to identify cell clusters. The first round of clustering analysis was performed on individual samples. We divided the genome into 5kb consecutive windows and then scored each cell for any insertions in these windows, generating a window by cell binary matrix for each sample. We filtered out those windows that are generally accessible in all cells for each sample using z-score threshold 1.65. Based on the filtered matrix, we then carried out dimension reduction followed by graph-based clustering to identify cell clusters. We called peaks for each cluster using the aggregated profile of accessibility and then merged the peaks from all clusters to generate a union peak list. Based on the peak list, we generated a cell-by-peak count matrix and used Scrublet (Wolock et al., 2019) to remove potential doublets. Next, to carry out the second round of clustering analysis, we merged peaks called from all samples to form a reference peak list. We then generated a single binary cell-by-peak matrix using cells from all samples and again performed the dimension reduction followed by graph-based clustering to obtain the final cell clusters across the entire dataset.

### Doublet removal

We applied Scrublet to the cell-by-peak count matrix with default parameters. Doublet scores returned by Scrublet were then used to fit a two-component Gaussian mixture model using the “BayesianGaussianMixture” function from the python package “scikit-learn”. The component with larger mean doublet score is presumably formed by doublets and cells belonging to it were removed from downstream analysis.

### Dimension reduction

To find the low-dimensional manifold of the single cell data, we adapted our previously published method, SnapATAC (Fang et al., 2020), to reduce the dimensionality of the peak by cell count matrix. The previous iteration of SnapATAC utilized spectral embedding for dimension reduction. To further increase the performance and scalability of spectral embedding, we applied the Nyström method (Bouneffouf and Birol, 2016) for handling large datasets. Specifically, we first randomly sampled 35,000 cells as the training data. We then computed the Jaccard index between each pair of cells in the training set and constructed the similarity matrix *S*. We computed the matrix *P* = *D*^−1^ *S*, where *D* is the diagonal matrix such that *D_ii_* = Σ*_j_ S_ij_*. The eigendecomposition was performed on *P* and the eigenvector with eigenvalue 1 was discarded. From the rest of the eigenvectors, we took the first 30 of them corresponding to the largest eigenvalues as the spectral embedding of the training data. We utilized the Nyström method to extend the embedding to the data outside the training set. Given a set of unseen samples, we computed the similarity matrix *S*′ between the new samples and the training set. The embedding of the new samples is given by *U*′ = *S*′*UΛ*^−1^, where *U* and *Λ* are the eigenvectors and eigenvalues of *P* obtained in the previous step.

### Correction of Batch Effects

Inspired by the mutual nearest neighbor batch-effect-correction method (Haghverdi et al., 2018), we developed a variant using mutual nearest centroids to iteratively correct for batch effects in multiple donor samples. Specifically, after dimension reduction we performed k-means clustering on individual replicate or donor sample with k equal to 20. We choose this number because the number of major clusters in a given tissue sample is typically less than 20. We then computed the centroid for each cluster and identified pairs of mutual nearest centroids across different batches. These mutual nearest centroids were used as the anchors to match the cells between different batches and correct for batch effects as described previously (Haghverdi et al., 2018). We found that the result can be further improved by performing above steps iteratively. However, too many iterations may lead to over-correction. We therefore used two iterations in this study.

### Graph-based clustering algorithm

We constructed the k-nearest neighbor graph (k-NNG) using low-dimensional embedding of the cells with k equal to 50. We then applied the Leiden algorithm (Traag et al., 2019) to find communities in the k-NNG corresponding to cell clusters. The Leiden algorithm can be configured to use different quality functions. The modularity model is a popular choice but it suffers from the issue of resolution-limit, particularly when the network is large (Traag et al., 2011). Therefore, we used the modularity model only in the first round of clustering analysis to identify initial clusters. In the final round of clustering, we chose the constant Potts model as the quality function since it is resolution-limit-free and is better suited for identifying rare populations in a large dataset (Traag et al., 2011). To determine the optimal number of clusters, we varied the resolution parameter in the Leiden algorithm and computed the clustering stability and diversity under each resolution. Cluster stability was defined as the consistency, measured by the average adjusted rand index, of results from five independent clustering analyses on perturbed inputs. The perturbation was introduced in a way that 2% of the edges were randomly selected and subjected to removal. To compute the cluster diversity, i.e., the extent to which different replicates are uniformly represented, we first grouped the cells based on their tissue of origin and then based on the experimental batch. We counted the cells for each combination and normalized by the total number of cells in the corresponding sample. For each tissue, normalized entropy was computed across batches. The average entropy across all tissues in the cluster were taken as the cluster diversity. Finally, we selected the highest resolution that had stability >0.9 and diversity >0.9.

### Iterative clustering analysis of major cell clusters

To further investigate the heterogeneity of identified cell clusters, we performed another round of clustering on 27 out of 54 cell clusters that had enough cells (> 1000) and minimal batch effects (diversity > 0.9), i.e., replicates are almost equally represented. For each of these cell clusters, we performed dimension reduction, batch correction and graph-based clustering as above. To avoid over-clustering, we selected the resolution parameter that lead to stable clustering results (stability > 0.9). 15 out of 27 cell clusters under investigation were found to contain more than one subcluster.

### Generating the union peak set

For each cluster, peak calling was performed on Tn5-corrected single-base insertions (each end of the Tn5-corrected fragments) using the MACS2 (Zhang et al., 2008) callpeak command with parameters “–shift -100 –extsize 200 –nomodel –call-summits –nolambda –keep-dup all -q 0.01”, filtered by the hg38 blacklist version 2 (downloaded from https://github.com/Boyle-Lab/Blacklist/tree/master/lists). To compile a union peak set, we combined peaks from all clusters and extended the peak summits by 250 bp on either side. Overlapping peaks were then handled using an iterative removal procedure. First, the most significant peak, *i.e.*, the peak with the smallest p-value, was kept and any peak that directly overlapped with it was removed. Then, this process was iterated to the next most significant peak and so on until all peaks were either kept or removed due to direct overlap with a more significant peak.

### Computing relative accessibility scores

We define an accessible locus as the minimal genomic region that can be bound and cut by the Tn5 enzyme. We use *L* ⊂ *N* to represent the set of all accessible loci. We further define a pseudo-locus as the set of accessible loci that relates to each other in certain meaningful way (for example, nearby loci, loci from different alleles). In this example, pseudo-loci correspond to peaks. We use {*d_i_* | *d_i_* ⊂ *L*} to represent the set of all pseudo-loci. Let *a_l_* be the accessibility of accessible locus *l*, where *l* ∈ *L*. We define the accessibility of pseudo-locus *d_i_* as *A_i_* Σ*_k∈d_i_a_k__*, *i.e.*, the sum of accessibility of accessible loci associated with di. Let *C_j_* be the library complexity (the number of distinct molecules in the library) of cell *j*. Assuming unbiased PCR amplification, then the probability of being sequenced for any fragment in the library is: 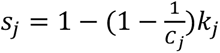 where *k_j_* is the total number of reads for cell *j*. If we assume that the probability of a fragment present in the library is proportional to its accessibility and the complexity of the library, then we can deduce that the probability of a given locus *l* in cell *j* being sequenced is: *p_ij_* ∝ *a_i_C_j_s_j_* . For any pseudo-locus *d_i_*, the number of reads in *d_i_* for cell *j* follows a Poisson binomial distribution, and its mean is *m_ij_* = Σ*_k∈dipkj_* ∝ *C_j_s_j_*Σ*_k∈d_i_a_k__* = *C_j_s_j_A_j_* . Given a pseudo-locus (or peak) by cell count matrix *O*, we have: Σ*_j_ O_ij_* = Σ*_j_ m_ij_*. Therefore, 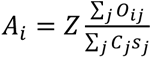, where *Z* is a normalization constant. When comparing across different samples the relative accessibility may be desirable as they sum up to a constant, *i.e.*, Σ*_i_ A_i_* = 1 x 10^6^. In this case, we can derive 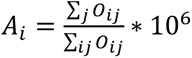

### Assigning cell types to cell clusters

To annotate the cell clusters, we first curated a set of marker genes from the PanglaoDB (Franzén et al., 2019) corresponding to expected cell types. We aggregated open chromatin fragments from each cluster and utilized the promoter accessibility, defined as RPM of +/- 1kb around TSS, as the proxy for gene activity. We then computed the raw cell type enrichment score as the logarithm of the geometric mean of marker genes’ activity. The final enrichment scores were obtained by applying two rounds of z-score transformation, first across cell types and then across cell clusters, on raw enrichment scores. For each cluster, we picked the cell type that showed strongest enrichment to make initial assignments. Finally, we manually reviewed these assignments and made adjustments based on focused consideration of marker gene accessibility in conjunction with information about tissue(s) of origin.

### Identification of cell type-restricted peaks

We used a Shannon entropy-based method (Schug et al., 2005) to identify cell type-specific peaks. Given the relative accessibility scores of a peak across clusters, we first converted the scores to probabilities: *p_i_* =*q_i_*/Σ*_i_ q_i_*. The entropy was then calculated by: *H_p_* = −Σ*_t_ p_t_* log_2_(*p_t_*). The specificity score is Q*_p_*_|*t*_ = *H_p_* – log_2_ (*p_t_*). To estimate the statistical significance of specificity scores, we assumed that under the null hypothesis each peak has an average accessibility level across all cell types and that the log base 2 of the cell-type-dependent fold changes from the average level follow a normal distribution with mean equal to zero and standard deviation *s*. The value of *s* was estimated using the top 50% least variable peaks, and 500,000 samples were then drawn to form the empirical distribution of *Q_p_* that are used to determine the p-values of specificity scores. The cell-type-restricted peaks were then identified using a FDR cutoff of 0.1%.

### Cell-type enrichment analysis of fine-mapped GTEx eQTLs

The fine-mapped eQTLs (GTEx Analysis V8) in each of the 25 tissues were downloaded from the GTEx portal (https://gtexportal.org). For each tissue, we first identified the overlapping cCREs with its eQTLs. We then calculated the average of log-transformed accessibility scores of these peaks in each of the 54 cell types. This yielded a tissue by cell-type table containing raw cell-type enrichment scores of eQTLs from each tissue. The raw enrichment scores were then normalized row-wise using z-score transformation. For each tissue, we defined the maximum cell-type enrichment as the largest value of z-scores across 54 cell types. In general, we found that homogenous tissues tend to have higher maximum cell-type enrichment than tissues that are more heterogenous.

### Differential peak analysis

To carry out differential peak analysis between foreground set and background set, we first removed all peaks with fold changes of relative accessibility less than 2. For each peak, we then built a full model and a reduced model.

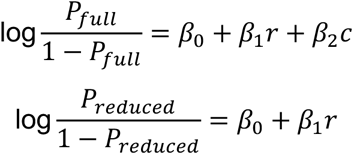

*P_reduced_*and *P_full_* represent the likelihood of the reduced model and full model respectively. *r* contains the logarithm of the number of fragments. *c* is a categorical variable indicating if the cell comes from the foreground or the background. We then used a likelihood ratio test framework to determine whether the full model provided a significantly better fit of the data than the reduced model. We selected the sites using a 5% FDR threshold (Benjamini-Hochberg method).

### Identification of fibroblast core signature and subtype-specific signatures

We first performed pairwise differential peak analysis for the six fibroblast subtypes. We then defined fibroblast core signature as peaks that are shared by all subtypes and were not called as differentially accessible in any of the pairwise comparison. Likewise, we defined the specific signature for a subtype as peaks that are differentially more accessible in the given subtype for every pairwise comparison.

### Measuring the similarity of chromatin accessibility profiles between cell types identified by sci-ATAC-seq and bulk biosamples

We downloaded bulk DNase-seq data from the ENCODE portal. We excluded samples collected at embryonic stage or originated from kidney, bladder or brain tissues, as we did not perform experiments on those tissues. As a result, 638 datasets were kept for downstream analysis. For each of the DNase-seq datasets, we calculated its Pearson correlation coefficient with 54 identified cell types based on RPKM values at identified cCREs. These correlation scores were then scaled using z-score transformation across 54 cell types. We used the maximum of scaled correlation scores to represent each biosample’s overall similarity with sci-ATAC-seq cell types.

### Identification of cCRE modules

A cCRE module is defined as co-accessible regions or regions that share similar accessibility pattern across cell types. We set a large k equal to 150 in k-mean clustering in order to capture complex patterning of 756,414 cCREs across 54 cell types. While the large number of clusters can better represent the complexity of the data, it also raises challenges for interpretability and downstream analysis. To address this, we further aggregated the 150 clusters into 51 super-clusters or CRE modules using hierarchical clustering. These 51 CRE modules were then retained for functional analysis and sequence motif analysis.

### Explaining cell-type specificity of CRE modules by deep learning

We used machine learning to investigate the extent to which the nucleotide sequences contribute to the cell type-specific chromatin accessibility pattern represented by the 51 cCRE modules. Specifically, we designed a sequence-to-module convolutional neural network (CNN) that uses one-hot-encoded DNA sequence (*A* = [1,0,0,0], *C* = [0,1,0,0], *G* = [0,0,1,0], *T* = [0,0,0,1]) as input to predict the module class for every cCRE. The architecture of CNN consists of a sequence of convolutional layers. Each convolutional layer has 64 filters with varying width. The first convolutional layer uses a filter width of 25 bp to scan the 500 bp region for relevant sequence motifs. This layer is then followed by 5 dilated convolutional layers (filter width 3) where the dilation rate doubles at every layer. A fully connected softmax layer is used after the convolutional layers to get module classes as the output. To ensure each module is uniformly represented in the training and testing datasets, we randomly selected 100 cCREs from each module to form the testing dataset. From the remaining cCREs, we then used oversampling to randomly sample 20,000 cCREs from each module to form the training dataset. We applied the Adam optimization algorithm to train the model until the validation accuracy stopped improving. To help interpret the model, we used the TF-MoDISco algorithm (Shrikumar et al., 2018) to extract the sequence motif features from the model and used TOMTOM (Gupta et al., 2007) to identify matched known TF motifs from a public database (Weirauch et al., 2014).

### Identification of candidate driver TFs

We used the Taiji pipeline (Zhang et al., 2019) to identify candidate driver TFs in each cell cluster. Briefly, for each cell type cluster, we constructed the TF regulatory network by scanning TF motifs at the accessible chromatin regions and linking them to the nearest genes. The network is directed with edges from TFs to target genes. The genes’ weights in the network were determined based on the relative accessibility of their promoters. The weights of the edges were calculated by the relative accessibility of the promoters of the source TFs. We then used the personalized PageRank algorithm to rank the TFs in the network.

### Comparing chromatin accessibility landscapes between adult and fetal cell types

To compare our dataset with the recent cell atlas of fetal chromatin accessibility (Domcke et al., 2020), we downloaded the bigwig files for different cell types in fetal tissues and converted the genomic coordinates from GRC37 (hg19) to GRCh38. In order to make a comparison, we focused on cell types present in eight organs that are profiled in both studies, including heart, intestine, muscle, adrenal gland, pancreas, lung, stomach, and liver. For each cell type, we then calculated the signal enrichment in the union peak list obtained by merging peaks from adult and fetal cell types. We applied quantile normalization to the resulting signal enrichment scores in order to mitigate technical or batch effects between the two datasets. We then compared the enrichment scores between adult and fetal cell types using Pearson correlation. To remove noise from the correlation calculation, for each pair of cell types we excluded regions that had enrichment scores less than 1 in both cell types from the calculation. To estimate the significance level of correlation scores, we used correlation scores from unmatched cell types to build a null model. We observed that these scores were roughly Gaussian distributed, and we used the sample mean and variance to parameterize a Gaussian model for computing p-values of correlation scores. To identify adult-specific peaks, for each peak we obtained the maximum value of enrichment scores across cell types in adult and fetal cell types respectively. We then log-transformed the maximum scores and computed the fold change between adult and fetus. We retained peaks with a fold change greater than 1.5 as adult-specific peaks. We used a similar strategy with some modifications when comparing the peaks in the same cell types from adult and fetus. Instead of taking the maximum, we compared average enrichment scores and used a more stringent cutoff of 2 for fold change thresholding.

### Generation of bigwig tracks

Each Tn5-corrected insertion was extended in both directions by 100 bp to form a 200-bp fragment. We then counted the number of fragments overlapping with each base on the genome and generated a bedgraph file. The bedgraph file was converted to bigwig file using the “bedGraphToBigWig” tool.

### Linking cCREs to target genes

We downloaded the chromosome interactions called from published promoter capture Hi-C data in 14 human tissues (Jung et al., 2019). In each tissue, we first filtered the chromosome interactions using a lenient p-value cutoff of 0.1. We then created the chromosome interaction matrix using the normalized interaction frequency. The interaction matrices from 14 tissues were then averaged to get the final interaction matrix. We applied the Activity-by-Contact (ABC) Model (Fulco et al., 2019) to compute the ABC Score for each cCRE-gene pair as the product of Activity (chromatin accessibility) and Contact (interaction frequency), normalized by the product of Activity and Contact for all other cCREs. We retained all distal cCRE-gene connections with an ABC score greater than 0.02.

### Estimating cell-type composition for tissues by deconvolution of bulk chromatin accessibility profiles

We selected 500 cCREs that were most specifically accessible in each of the 54 cell types according to the specificity scores defined above. These cCREs were used to create a signature cCRE matrix, which contained accessibility scores of 19,591 distinct cCREs across 54 cell types. To estimate the fractions of 54 cell types from chromatin accessibility profiles of bulk tissue samples, we solve the linear equation: *Sb*= *v*, where *S* is the cell-type by cCRE signature matrix, *b* is a column vector containing fractions of 54 cell-types, and *v* is the bulk chromatin accessibility scores of 19,591 signature cCREs. We applied two different algorithms, non-negative least squares (NNLS) and support vector regression (SVR), for solving the equations. We found that the two algorithms show comparable performance while SVR performs a little better than NNLS.

### GWAS variant enrichment

We used linkage disequilibrium (LD) score regression (Bulik-Sullivan et al., 2015) v1.0.1 to estimate genome-wide GWAS enrichment for disease and non-disease phenotypes within cell type resolved cCREs (peaks called on each cell cluster via MACS2 (Zhang et al., 2008) using the above parameters). We compiled published GWAS summary statistics for complex diseases (Bentham et al., 2015; Bronson et al., 2016; Consortium, 2019; Cordell et al., 2015; Jansen et al., 2019; Ji et al., 2017; Jin et al., 2016; Luo et al., 2017b; Mahajan et al., 2018; Malik et al., 2018; Michailidou et al., 2017; Nielsen et al., 2018; Nikpay et al., 2015; Okada et al., 2014; Paternoster et al., 2015; Pividori et al., 2019; Sakornsakolpat et al., 2019; Schafmayer et al., 2019; Shadrina et al., 2019; Tachmazidou et al., 2019; Tin et al., 2019; Watanabe et al., 2019; Wiberg et al., 2019; Wuttke et al., 2019) and endophenotypes (Astle et al., 2016; Hoffmann et al., 2018; Kemp et al., 2017; Kilpeläinen et al., 2016; Manning et al., 2012; Saxena et al., 2010; Shrine et al., 2019; Strawbridge et al., 2011; Teumer et al., 2018; Warrington et al., 2019) within European populations. Using cell type resolved cCREs as a binary annotation, we created custom partitioned LD score files by following the steps outlined in the LD score estimation tutorial. As background annotations, we included all baseline annotations in the baseline-LD model v2.2 as well as partitioned LD scores created from all merged cCREs. For each trait, we used LD score regression to then estimate coefficient z-scores for each cell type relative to the background annotations. We used the coefficient z-scores to compute one-sided p-values and used the Benjamini-Hochberg procedure to correct for multiple tests.

### Fine mapping

We performed genetic fine mapping for GWAS of diseases and endophenotypes that had sufficient coverage (i.e., were at least imputed into 1000 Genomes). For GWAS with available fine mapping data, we took 99% credible sets directly from the supplemental tables. For GWAS without available fine mapping data, we calculated approximate Bayes factors (Wakefield, 2009) (ABF) for each variant assuming prior variance ω = 0.04. For every trait, we obtained index variants for each locus from the supplemental tables of the respective study. We extracted all variants in at least low linkage disequilibrium (r2 > 0.1 using the European subset of 1000 Genomes Phase 3 (Auton et al., 2015)) in a large window (±2.5 Mb) around each index variant. We calculated posterior probabilities of association (PPA) for each variant by dividing its ABF by the cumulative ABF for all variants within the locus. We then defined 99% credible sets for each locus by sorting variants by descending PPA and keeping variants adding up to a cumulative PPA of 0.99.

### Predicting the effects of non-coding variants on TF binding

To identify SNPs that affect TF binding, we employed deltaSVM models as described previously (Yan et al., 2021). Briefly, 40 bp sequences centered on each SNP were used as input to 94 previously trained and validated TF models. For each SNP, we predicted the binding scores for both alleles by running “gkmpredict”. A SNP is considered to be bound if the binding score passes the pre-defined threshold for either allele. Among those SNPs, deltaSVM scores were calculated using the “deltasvm.pl” script and SNPs with deltaSVM scores passing the threshold for the corresponding model are predicted to affect TF binding.

### DATA AND SOFTWARE AVAILABILITY

The GEO accession number for the sequencing data and processed data files in this paper is GSE165659.

